# Neuron-specific cTag-CLIP reveals cell-specific diversity of functional RNA regulation in the brain

**DOI:** 10.1101/244905

**Authors:** Yuhki Saito, Yuan Yuan, Ilana Zucker-Scharff, John J. Fak, Yoko Tajima, Donny D. Licatalosi, Robert B. Darnell

**Affiliations:** Laboratory of Molecular Neuro-oncology and Howard Hughes Medical Institute, The Rockefeller University, 1230 York Avenue, New York, New York 10065, USA.; Center for RNA Science and Therapeutics, Case Western Reserve University, Cleveland, Ohio 44106, USA.

**Keywords:** CLIP, alternative splicing, intron retention, cTag-CLIP, RNA-binding protein, NOVA, single cell RNA regulation

## Abstract

RNA-binding proteins (RBPs) regulate genetic diversity, but the degree to which they do so in individual cell-types *in vivo* is unknown. We employed NOVA2 cTag-CLIP to generate functional RBP-RNA maps from single neuronal populations in the mouse brain. Combining cell-type specific data from *Nova2-cTag* and *Nova2* conditional knock-out mice revealed differential NOVA2 regulatory actions (e.g. alternative splicing) on the same transcripts in different neurons, including in cerebellar Purkinje cells, where NOVA2 acts as an essential factor for proper motor coordination and synapse formation. This also led to the discovery of a mechanism by which NOVA2 action leads to different outcomes in different cells on the same transcripts: NOVA2 is able to regulate retained introns, which subsequently serve as scaffolds for another *trans-acting* splicing factor, PTBP2. Our results describe differential roles and mechanisms by which RBPs mediate RNA diversity in different neurons and consequent functional outcomes within the brain.

## INTRODUCTION

Alternative splicing (AS) plays spectacular roles in mammalian biology that are especially evident in the brain, where it generates complexity from individual transcripts from a limited genome to act as a major driver of system complexity (Darnell, 2013; Raj and Blencowe, 2015; Zheng and Black, 2013). The first example came from Amara and Evans, who discovered that the gene for calcitonin is alternatively spliced in the brain to generate the neurotransmitter peptide, CGRP (Amara et al., 1982). Subsequently, enormous diversity was discovered in *Drosophila* where stochastic AS generates mutually exclusive isoforms of DSCAM (Schmucker et al., 2000; Zipursky and Grueber, 2013) that serve to regulate appropriate axon fasciculation, with a parallel system in mammalian protocadherins, which use analogous stochastic mechanisms to regulate mammalian axonal outgrowth (Chen et al., 2012; 2017; Mountoufaris et al., 2017; Wu and Maniatis, 1999). Other examples in mammalian brain development have included documentation of AS switching controls by neuronal proteins such as PTBP2 (Coutinho-Mansfield et al., 2007; Li et al., 2014; Licatalosi et al., 2012; Makeyev et al., 2007), NOVA (Eom et al., 2013; Jensen et al., 2000; Ruggiu et al., 2009; Saito et al., 2016; Ule et al., 2005b; Yano et al., 2010), RBFOX (Gehman et al., 2012; Lovci et al., 2013; Weyn-Vanhentenryck et al., 2014), MBNL (Charizanis et al., 2012; Wang et al., 2012), nElavl (Ince-Dunn et al., 2012), and nSR100 (Calarco et al., 2009; Quesnel-Vallières et al., 2015) that govern proper early neuronal development.

An unresolved question in these studies has been what role such factors play in cell-type specific identity, development and function within the brain. Such issues have been challenging because dissociation and purification of individual cell types, for example by laser capture microdissection or antibody-mediated panning methods can produce limited numbers of cells and/or neurons, astrocytes or microglia missing their processes, which may account for large percentages of the volumes of such cells. Moreover, while such methods have been used for RNA sequencing, they are not informative about RBPs-mediated cell-specific RNA regulation. We developed a means for purifying native RNA-protein regulatory complexes from intact flash-frozen brains termed CLIP (Licatalosi et al., 2008; Ule et al., 2003). We recently modified CLIP using a technology termed cTag-PAPERCLIP to generate a conditionally AcGFP-tagged poly(A) binding protein PABPC1 (Hwang et al., 2017); after isolation of brain, CLIP and AcGFP purification, PAPERCLIP was able to differentiate polyadenylated (poly(A)) RNAs that were unique to excitatory and inhibitory neurons, and distinguish these from microglia and astrocytes. This led to identification of microglia-specific poly(A) changes after inflammatory brain stimulation (LPS) (Hwang et al., 2017) that were otherwise undetectable using standard methods to purify brain microglia.

Here we explore whether cTag-CLIP can be applied more generally for purification of a subset of RNA-protein regulatory interactions in specific cell-types. We developed cTag-CLIP to purify NOVA2, a neuron-specific RNA regulatory protein together with its bound RNA targets. NOVA1 (Buckanovich and Darnell, 1997) and NOVA2 (Yang et al., 1998) proteins have been implicated in excitatory-inhibitory neuronal balance, from their original identification as targets in paraneoplastic opsoclonus-myoclonus-ataxia (POMA), and from studies in mice. In the latter, *Nova2* haploinsufficiency leads to spontaneous epilepsy (Eom et al., 2013), and homozygous loss of either or both NOVA isoforms leads to early postnatal death and developmental defects in neuronal migration and axonal guidance (Jensen et al., 2000; Ruggiu et al., 2009; Saito et al., 2016; Yano et al., 2010). Some aspects of these phenotypes relate to the role NOVA plays in AS; NOVA regulates the *agrin* Z^+^ alternative exon, shown by Sanes and colleagues (Burgess et al., 1999) to be essential for acetylcholine clustering at the neuromuscular junction (NMJ; Gautam et al., 1996). In *Nova1/2* null mice acetylcholine clustering and synaptic transmission at the NMJ can be rescued by a constitutive *agrin* Z+ transgene (Ruggiu et al., 2009), although mice remain paralyzed, pointing to additional unknown neuronal roles for NOVA proteins.

In this study, we harnessed cTag technology, in which AcGFP-tagged RBPs are conditionally expressed using the Cre/Lox system (Hwang et al., 2017), to access NOVA2 *in vivo* at cell-type specific resolution. NOVA2 cTag-CLIP using high-affinity anti-GFP antibody mixtures allows purification of NOVA2-RNA complexes to generate maps of functional NOVA2-RNA interaction from Cre-expressing cells. Integrating data from NOVA2 cTag-CLIP with RNA-seq in *Nova2*-conditional knock out (Nova2-cKO) mice in different neuronal types led us to discover distinct NOVA2 properties and functions on the same transcripts in different cellular contexts, including in cerebellar Purkinje cells, where it acts as an essential factor for proper motor coordination and synapse formation. Exploring these findings further led to the discovery of a mechanism by which NOVA2 action leads to different outcomes in different cells on the same transcripts: NOVA2 acts as a *cis*-acting AS element to regulate intron retention (IR), which subsequently serve as cell-specific scaffolds with which to alter RBP-RNA dynamic interactions of another *trans-acting* AS factor, PTBP2. We discuss this new complexity in understanding how RBPs mediate RNA diversity and neuron-type specific functional biology within the brain.

## RESULTS

### Alternative splicing and RBPs expression diversity across the CNS regions and neuronal types

To compare our strategy with what is currently known about AS diversity across CNS regions, we re-analyzed publically available RNA-seq datasets of multiple CNS regions and ages with the Quantas pipeline (Charizanis et al., 2012; Yan et al., 2015). Over 3,000 cassette-type AS events could be identified as showing significant differences (|*Δ*exon inclusion rate| (|*Δ*I|)>=0.1 and FDR<0.05) during mouse cortical development and among the adult CNS regions (cerebral cortex, cerebellum, and spinal cord), and over 1,800 AS events were recognized between the neonatal cortex and the mixture of mid, hindbrain, and cerebellum (Figure 1A, S1A, and S1B). The AS events regulated by neural-enriched splicing factors, including NOVA, nSR100, PTBP2, MBNL, and RBFOX proteins, were determined by re-analyzing the publically available RNA-seq datasets of each RBP-KO mouse, with total AS event numbers changed in each RBP-KO mouse brain shown in Figure S1C and Table S1. Investigating the relative contribution of neuronal-enriched RBPs to spawn this AS diversity across the CNS regions and during mouse cortical development revealed that NOVA2 was the top ranked splicing regulator providing AS diversity across CNS regions (Figure 1A).

**Figure 1.**
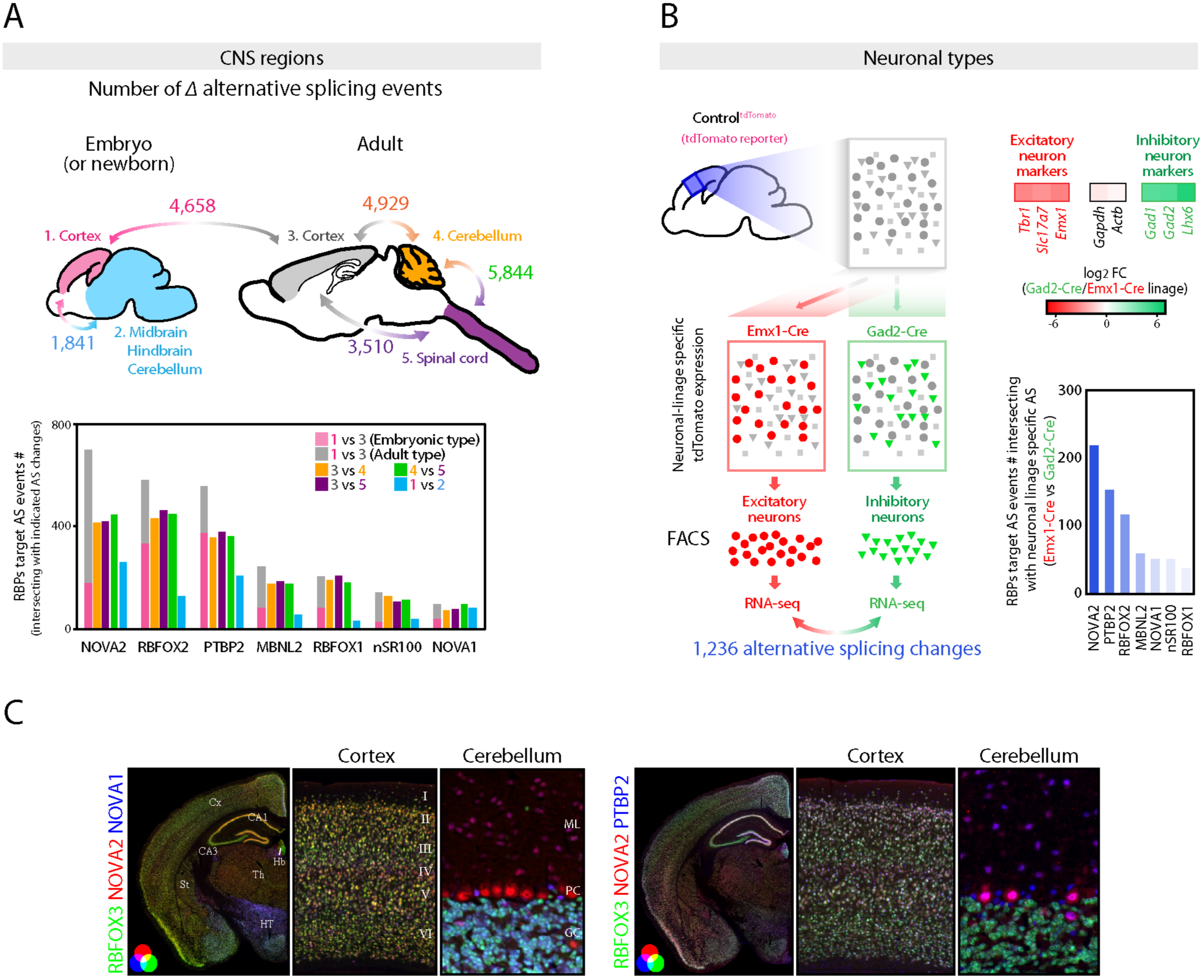
Diversity of alternative splicing and RNA binding proteins expression within the CNS regions and neuronal types. (A) AS diversity across the CNS regions and during cortical development. *Upper panel* shows the schematics illustrating the summary of AS changes determined by RNA-seq data across the mouse CNS regions and during mouse cortical development. The number indicates the significantly changed cassette-type AS event number (FDR<0.05, |*Δ*I|>=0.1) between two tissues indicated by bidirectional arrows. *Lower graph* shows the AS event numbers intersecting with the AS events changing in each RBP-KO mouse brain (FDR<0.05, |*Δ*I|>=0.1). (B) AS diversity between neuronal-types. Schematics illustrating the strategy for AS analysis between two distinct neuronal types prepared from e18.5 cortex of ^tdTomato^ reporter mouse which expresses ^tdTomato^ in Cre dependent fashion (*Left panel*). 1,236 AS events were significantly changed between two neuronal types (FDR<0.05, |*Δ*I|>=0.1). Heatmaps show the excitatory or inhibitory neuron markers enrichment determined by RNA-seq (*Upper right*). AS event numbers intersecting with AS events changing in each RBP-KO mouse brain (FDR<0.05, |*Δ*I|>=0.1) (*Lower right*). (C) RBPs expression diversity within the CNS regions and neuronal types. NOVA2, NOVA1 and RBFOX3 or PTBP2 immunofluorescence staining images in 4 weeks old mouse brain sections. “See also Figure S1 and Table S1.”

Neurons and glia, the main cell-types in the brain, use different sets of AS exons (Zhang et al., 2014). It is unclear, however, the extent to which AS diversity exists among individual neuronal subtypes, nor what might govern such regulation. As a baseline for demarcating neuronal type regulatory factors that might govern cell-specific AS events, single cell suspensions prepared from embryonic day 18.5 (e18.5) cortex of ^tdTomato^ reporter mice (Control^tdTomato^) crossed with Gad2-Cre (inhibitory neuron-linage) or Emx1-Cre (cortical excitatory neuron-linage) driver mouse (Control^tdTomato,Gad2-Cre^ or Control^tdTomato,Emx1-Cre^, respectively) were subjected to FACS and RNA-seq analysis (Figure 1B). Cre-dependent ^tdTomato^ expression was confirmed by immunofluorescence staining for ^tdTomato^ (Figure S1D), and as anticipated, GABAergic inhibitory neuronal markers (e.g. *Gad1, Gad2*, and *Lhx6*) or glutaminergic excitatory neuronal markers (e.g. *Tbr1, Slc17a7*, and *Emx1*) were significantly enriched in inhibitory neuronal-linage or excitatory neuronal-linage, respectively (Figure 1B). Cell-type specific expression of levels of AS regulators were then assessed and found to be non-uniform, such that *Nova1, Mbnl1, Elavl2, Elavl4*, and *Celf4* were significantly enriched in inhibitory neuronal-linage, and *Nova2, Rbfox1, Rbfox3, Celf2*, and *Celf6* tended to be enriched in excitatory neuronal-linage (Figure S1E). The comparison of exon inclusion rate between inhibitory and excitatory neuronal-linage identified 1,236 significant AS changes (FDR<0.05, |*Δ*I|>=0.1) (Figure 1B, S1F, and Table S1), over two hundred of which overlapped with the NOVA2 target AS events (Figure 1B and S1C). Neuronal-enriched *trans-acting* splicing regulators shows diverse expression patterns across the CNS regions and neuronal types (Figure 1C), suggesting that the identified AS diversity across the CNS regions and neuronal-types are spawned by the expression and action of individual trans-acting splicing factors such as NOVA2.

### Discovery of neuronal linage selective NOVA2 AS targets with new mouse models

NOVA was the first RBP whose *in vivo* RNA-binding map was generated with CLIP (Ule et al., 2003), and the subsequent application of high throughput sequencing to CLIP (HITS-CLIP; (Licatalosi et al., 2008)) demonstrated the ability of the method to generate robust transcriptome-wide regulatory protein-RNA interaction maps *in vivo*. To generate NOVA2 CLIP maps with single cell-type specific resolution in intact CNS tissue, we generated “Nova2-cTag” mice that conditionally express AcGFP-tagged NOVA2 (NOVA2-AcGFP) in a Cre-dependent manner (Figure 2A and 2B). A knock-in strategy was employed, targeting the endogenous *Nova2* locus, in order to maintain wild-type *Nova2* RNA stoichiometry and alternative processing (Figure S2A). Nova2-cTag heterozygous and homozygous mice had no apparent phenotype.

**Figure 2.**
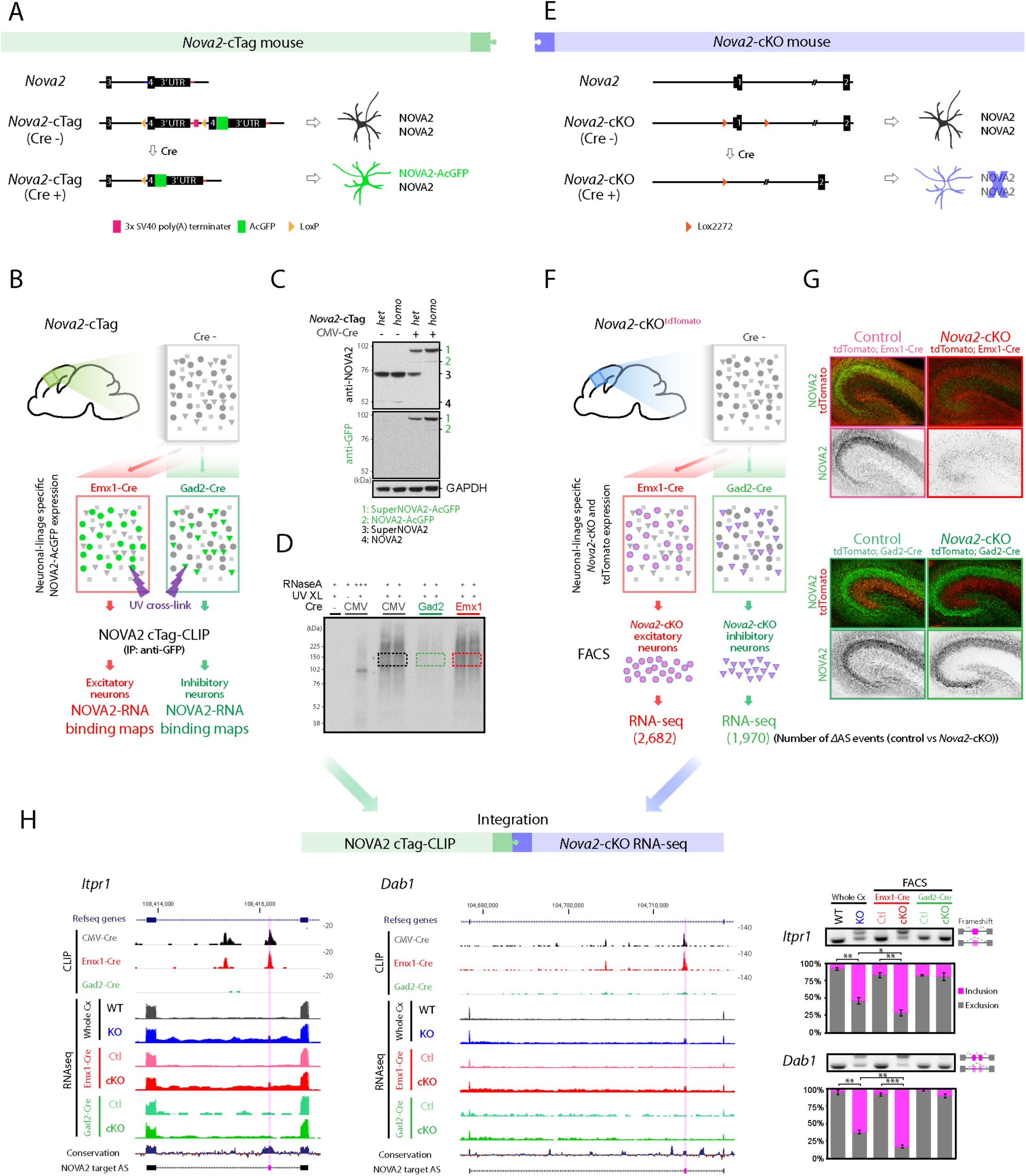
Generation of Nova2-cTag and *Nova2*-cKO mouse models discovering neuronal linage selective NOVA2 AS targets. (A) Schematics illustrating simplified Nova2-cTag mouse model providing selective AcGFP-tagged NOVA2 expression in specified neuronal-type *in vivo*. (B) Schematics illustrating the strategy for selective NOVA2 cTag-CLIP in the targeted neuronal type *in vivo*. (C) Cre-dependent NOVA2-AcGFP protein expression. Indicated ouse brain lysate were subjected to immunoblot analysis with anti-NOVA2 or anti-GFP antibody. (D) Autoradiograph image of neuronal linage specific NOVA2 cTag-CLIP. *Colored box* regions were subjected to CLIP library cloning steps. (E) Schematics illustrating the *Nova2*-cKO model providing NOVA2 depletion from selective neuronal type *in vivo*. (F) Schematics illustrating the strategy for AS analysis in specific neuronal type (FDR<0.05, |*Δ*I|>=0.1, n=3). (G) NOVA2 depletion from selective neuronal populations. NOVA2 (*green*) and ^tdTomato^ (*red*) immunofluorescent staining images in the e18.5 hippocampus of indicated 4 mouse lines. (H) UCSC genome browser views of neuronal linage selective NOVA2 mediated AS events (*left panels*). RT-PCR confirmation of AS changes detected by RNA-seq (*right panels*) (n=3). *; p<0.05, **; p<0.01, ***; p<0.001. “See also Figure S2 and Table S1.”

To examine the RNA targets of AcGFP-tagged NOVA2 relative to endogenous NOVA2, we bred *Nova2*-cTag mice to a near ubiquitous Cre driver mouse (CMV-Cre) in mouse neurons (*Nova2*-cTag^CMV-Cre^). Immunoblot and immunofluorescent staining experiments on *Nova2*-cTag^CMV-Cre^ demonstrated successful and complete Cre-dependent expression of NOVA2-AcGFP (Figure 2C and S2B) at e18.5. We then undertook a side-by-side comparison of the RNA binding properties between native NOVA2 and AcGFP-tagged NOVA2. We undertook NOVA2-AcGFP HITS-CLIP (referred as NOVA2 cTag-CLIP) in the e18.5 cortex of Nova2-cTag^CMV-Cre^ mice, using a mixture of two high-affinity monoclonal anti-GFP antibodies (Figure 2D). Comparison of the results from 3 biologic replicate NOVA2 cTag^CMV^'^Cre^-CLIP experiments and native revealed NOVA2 binding to nearly identical (R=0.99) target transcripts with extremely similar CLIP peaks levels (R= 0.87, respectively) (Figure S2C) and NOVA2 binding to its known (Buckanovich and Darnell, 1997; Lewis et al., 1999) YCAY motifs in CLIP peaks (Figure S2E). Taken together, this data indicates that AcGFP-tagged NOVA2 is not detectably altered in its RNA binding properties *in vivo*.

We next applied NOVA2 cTag-CLIP to generate NOVA2-RNA interaction maps in two major neuronal-types: GABAergic inhibitory neurons and glutamatergic excitatory neurons within e18.5 cortex (n=3). *Nova2*-cTag mice were crossed with the cell-specific Cre drivers used above (*Nova2*-cTag^Gad2-Cre^ and *Nova2*-cTag^Emx1-Cre^, respectively). GFP immunofluorescent staining on these mice showed successful Cre-dependent expression of NOVA2-AcGFP in the appropriate subsets of neurons (Figure S2B). Comparing the NOVA2 cTag-CLIP tag number per transcript or per CLIP peaks showed significant enrichment of 923 and 791 transcripts and of 823 and 752 CLIP peaks in inhibitory (NOVA2-cTag^Gad2-Cre^) or excitatory (NOVA2-cTag^Emx1-Cre^) neurons, respectively (FDR<0.05, |log2 fold change|>=1). These results indicate that NOVA2 cTag-CLIP is able to discriminate large distinctions in NOVA2-RNA binding profiles between inhibitory and excitatory neurons (Figure S2D), and more, when taken together with prior cTagPAPERCLIP studies of PABPC1 (Hwang et al., 2017), suggest the general utility of the cTag-CLIP strategy to define RNA-protein regulation with cell-type specific resolution *in vivo*.

To search for NOVA2 directly-mediated AS events in selective neuronal populations, we first tried to define all AS events in *Nova2*-cKO (Figure 2E and 2F). Newly generated *Nova2-* cKO (Figure 2E and S2F) were crossed with the cell-specific Cre drivers used above, and Cre-dependent NOVA2 protein depletion from a selective neuronal population at e18.5 in *Nova2-* cKO mice was confirmed by immunofluorescent staining on 4 mouse lines (Control^tdTomato;Gad2-Cre^, *Nova2*-cKO^tdTomato;Gad2-Cre^, Control^tdTomato;Emx1-Cre^, and *Nova2*-cKO^tdTomato;Emx1-Cre^; Figure 2G and S2G). Control or *Nova2*-cKO ^tdTomato^ positive cells expressed in inhibitory or excitatory neurons were collected by FACS, and RNA-seq datasets prepared from each were analyzed for AS (Figure 2F and S2H). When the exon inclusion rate of RNA transcripts in control and *Nova2*-cKO inhibitory and excitatory neurons were compared, 1,970 or 2,682 AS events could be discriminated, respectively (Figure 2F and S2H). As examples (Figure 2H), we show NOVA2-dependent selective splicing in excitatory but not inhibitory neurons of a novel putative occult NMD *Itpr1* exon, and the Dab1.7bc exons where were previously identified as a NOVA2-dependent switch of Reelin-Dab1 signaling in cortex. The current approach was able to extend observations of NOVA2-dependent Dab1.7bc exons to implicate a specific to subset of (excitatory) cortical neurons (Yano et al., 2010). Importantly, this cell-type selective NOVA2-dependent AS was also found together with significantly increased NOVA2 cTag-CLIP peak height (as well as intron retention (IR; see below, Figure 5 and Figure 6)) in excitatory (Emx1-Cre) but not inhibitory (Gad2-Cre) neurons (Figure 2H), and these splicing changes determined by RNA-seq AS analysis were confirmed by RT-PCR (Figure 2H). Interestingly, *Nova2-* cKO^Emx1-Cre^ mice died between three and four weeks old and showed disorganized hippocampal CA1 and CA3 laminar structure (Figure S2I), which requires proper Reelin signaling (Caviness and Sidman, 1973). In addition to laminar disorganization of hippocampal excitatory neurons, the reduced thickness of CA1 stratum lacunosum-moleculare (SLM) and of dentate gyrus molecular layer was observed in 3 weeks old *Nova2*-cKO^Emx1-Cre^ mice (Figure S2J). These phenotypes in Nova2-cKO^Emx1-Cre^ were not observed in Nova2-cKO^Gad2-Cre^ mice (data not shown). We conclude that combining data from Nova2-cTag and *Nova2*-cKO mouse brain provides a powerful means of defining cell specific RNA regulation able to reveal new insight into brain function development, here indicating neuronal-linage selective NOVA2-mediated RNA regulation required for proper development of distinct neuronal subtypes and the structures they generate.

### NOVA2 characterizes cell-type specific exon skipping in inhibitory cerebellar Purkinje cells

NOVA2 is highly expressed in cerebellar GABAergic Purkinje cells (PCs), in contrast to relatively higher expression in forebrain glutamatergic excitatory neurons (Figure 1C). However, NOVA2 functions in adult PCs have been unexplored because *Nova2*-KO mice die 2–3 weeks after birth (Saito et al., 2016). Here we investigated NOVA2-mediated RNA regulatory networks in adult PCs by analysis of *Nova2*-cTag and *Nova2*-cKO mice models. *Nova2*-cTag mice were bred with three Cre driver mouse lines to generate mouse lines expressing NOVA2-AcGFP in the entire cerebellum or specifically in PCs versus granule cells (GCs) (using CMV, Pcp2, or vGlut1-Cre drivers, respectively; Figure 3A and S3A). NOVA2-AcGFP immunofluorescent staining showed highly restricted and anticipated expression patterns in cerebellum for each Cre driver mouse (Figure S3A). Comparison of pan-NOVA2 cTag-CLIP in 4 weeks old mouse cerebellum (NOVA2 cTag^CMV-Cre^-CLIP) and native NOVA2 CLIP revealed highly similar tags per transcript and per CLIP peak (Figure S3B), analogous to what was observed in cortex, indicating that the cTag strategy did not alter the RNA binding properties of NOVA2 in adult cerebellum.

**Figure 3.**
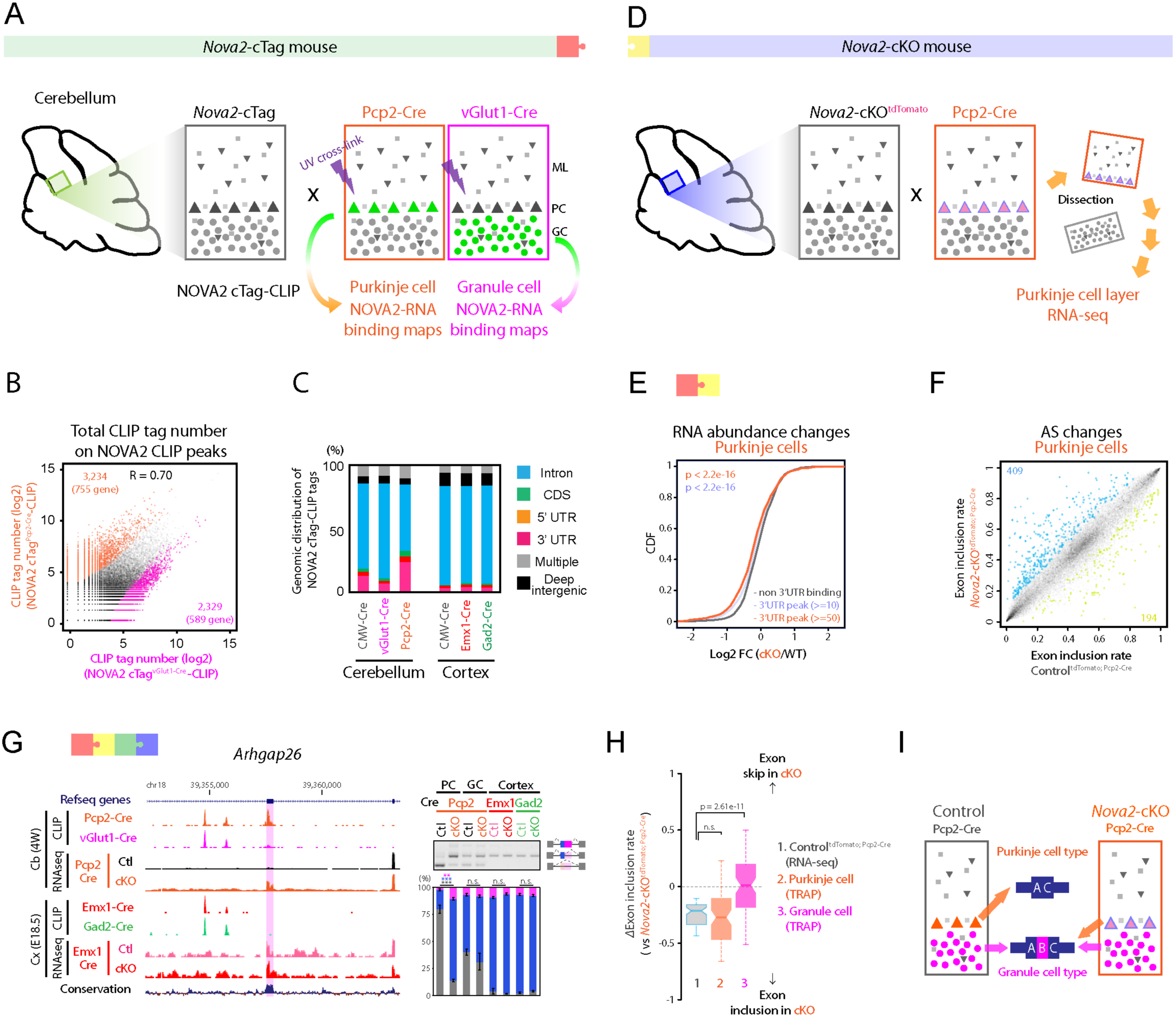
NOVA2 characterizes Purkinje-type exon skip in cerebellar Purkinje cells. (A) Schematics illustrating the overview for neuronal type specific NOVA2 cTag-CLIP in cerebellum. (B) The comparison of neuronal-type specific NOVA2 cTag-CLIP between Purkinje cells (PCs) and granule cells (GCs). Scatter plot shows the correlation of read counts on CLIP peaks between NOVA2 cTag^Pcp2-Cre^-CLIP to NOVA2 cTag^vGlut1-Cre^-CLIP. Each *black* dot represents a comparable CLIP peak between two neuronal-type. Each *orange* or *magenta* dot represents a significantly enriched CLIP peak in either NOVA2 cTag^Pcp2-Cre^-CLIP or cTag^vGlut1-Cre^-CLIP, respectively (FDR<0.05, |log2 FC|>=1). R: correlation coefficient. (C) Genomic distribution of NOVA2 and NOVA2 cTag-CLIP unique tags. (D) Schematics illustrating the strategy for AS analysis in PCs. (E) Transcript abundance changes upon NOVA2 depletion from PCs determined by RNA-seq. The empirical cumulative distribution function of NOVA2 targets binned by NOVA2 cTag^Pcp2-Cre^-CLIP peak tags. (F) AS changes depending on NOVA2 depletion from PCs. Scatter plot shows the exon inclusion rate of the 4 weeks old Control^tdTomato; Pcp2-Cre^ and Nova2-cKO^tdTomato; Pcp2-Cre^ determined by RNA-seq. Each *black* dot represents a comparative AS event. Each *blue* or *limegreen* dot indicates the significantly changed AS event which skipped in Nova2-cKO^tdTomato; Pcp2-Cre^ (409 AS events) or included in Nova2-cKO^tdTomato; Pcp2-Cre^ (194 AS events), respectively (n=3, FDR<0.05, |*Δ*I|>=0.1). (G) Discovery of PCs selective NOVA2 mediated AS event. UCSC genome browser view of around *Arhgap26* AS exon 11 (NM_175164) (*left*). RT-PCR confirmation of AS change detected by RNA-seq (*right panels*) (n=3). **; p<0.01, ***; p<0.001. (H & I) GCs-type exon inclusion in PCs of *Nova2*-cKO. ΔExon inclusion rate (ΔI) of 1; Control^tdTomato; Pcp2-Cre^ PCs layer (RNA-seq), 2; wild-type PCs (TRAP) or 3; wild-type GCs (TRAP; Mellén et al., 2012) versus Nova2-cKO^tdTomato; Pcp2-Cre^ PCs layer (RNA-seq) (H). Schematics illustrating the AS patterns switch from inhibitory PCs-type to excitatory GCs-type in *Nova2*-cKO^tdTomato; Pcp2-Cre^ PCs (I). “See also Figure S3 and Table S1.”

Comparison of the NOVA2 cTag-CLIP results in inhibitory Purkinje and excitatory granule neurons showed remarkable differences (Figure 3B and S3C). Large numbers of NOVA2 binding sites were significantly enriched in PCs versus GCs, either from analysis of NOVA2-cTag CLIP tags (4,840 versus 3,850 transcripts respectively; FDR<0.05, |log_2_ fold change|>=1); Figure S3C) or CLIP peaks (3,234 versus 2,329 peaks, respectively; FDR<0.05, |log_2_ fold change|>=1) (Figure 3B). The genomic distribution of CLIP peaks revealed NOVA2 in PCs preferentially binds the 3’UTR (25%) relative to other neuronal-types (less than 10% each; Figure 3C). Nonetheless, NOVA2 retained its YCAY motif binding enrichment across all examined NOVA2 cTag-CLIP peaks (Figure S2E and S3D). These results indicate that NOVA2 recognizes its direct YCAY binding targets independent of specific neuron-types, but for reasons that likely involves cell-specific interactions with other RBPs (see below and Discussion), discriminates among the same transcripts expressed in different neuronal types.

To investigate NOVA2-mediated regulation of RNA metabolism in individual cell types, including AS and transcriptome abundance, we next manually dissected PC enriched layers from acute cerebellar slices of Control^tdTomato; Pcp2-Cre^ and *Nora2*-cKO^tdTomato; Pcp2-Cre^ mice (Figure 3D and S3E), and subjected extracted RNA to RNA-seq analysis (n=3). This revealed significant enrichment of established PC markers (*Itpr1, Grid2, Calb1, Grm1*) in the dissected PC layer, relative to whole cerebellar and GCs enriched markers (*Neurod1, Slc17a7, Calb2, Rbfox3*), which were significantly depleted (FDR<0.01; Figure S3F). PC-specific enrichment of NOVA2 cTag-CLIP tags in 3’UTR prompted us to examine the NOVA2 dependent RNA abundance change in *Nova2-*cKO^tdTomato; Pcp2-Cre^ versus Control^tdTomato; Pcp2-Cre^ PCs, revealing significantly reduced levels of 3’UTR NOVA2 target transcripts in the absence of *Nova2* (Figure 3E and S3G). These observations suggest NOVA2 binding to 3’UTR mediates increased mRNA abundance regulation in cell-specific manner in Purkinje neurons.

We examined NOVA2-dependent AS changes in RNA-seq from the dissected PCs layer of *Nova2* Control^tdTomato;Pcp2-Cre^ and Nova2-cKO^tdTomato;Pcp2-Cre^ mice, revealing a total of 603 NOVA2-dependent AS changes (Figure 3F). Overlapping cerebellar and cortical NOVA2 cTag-CLIP and corresponding *Nova2-*cKO RNA-seq datasets revealed 396 Purkinje neuron selective NOVA2 target AS events (Figure S3H). For example, *Arhgap26*, the target antigen in a patient with subacute autoimmune cerebellar ataxia (Jarius et al., 2010), harbored an exon (exon11) that was NOVA2-regulated in a PC-restricted manner in *Nova2*-cKO^tdTomato;Pcp2-Cre^ and had a selective NOVA2 cTag^Pcp2-Cre^-CLIP peak closely positioned next to it (Figure 3G). RT-PCR result supported a NOVA2-dependent PC layer selective *Arhgap26* AS change (Figure 3G). NOVA2 mediated *Arhgap26* exon11 skipping in wild-type PCs (Control^tdTomato;Pcp2-Cre^), as evident by aberrant exon11 splicing inclusion in PCs layer of Nova2-cKO^tdTomato;Pcp2-Cre^ in a manner that was similar to the exon11 splicing inclusion pattern of control GCs (lane 2 and 3 in Figure 3G right panel).

To assess more generally whether NOVA2 was switching from GCs-like splicing to a unique PCs profile, we re-analyzed publically available translating ribosome affinity purification (TRAP) datasets of PCs and GCs with our AS analysis pipeline and identified a total of 6,163 AS events that were significantly changed between PCs and GCs (Figure S3I; n=4, FDR<0.05, |*Δ*I|>=0.1). 75 of 189 cassette-type NOVA2 target AS events in PCs layer in which NOVA2 significantly promoted exon skipping were compared in PCs and GCs. Comparing the average exon usage (*Δ*I) in wild type PCs, GCs and *Nova2*-cKO PCs revealed that the average *Δ*I of GCs exon inclusion was significantly increased (Figure 3H). These results are consistent with NOVA2 changing a default GCs type splicing profile mediating exon inclusion to a PC exon skipping pattern, and that in *Nova2* deficient PCs this AS pattern reverts to the GCs-type pattern (Figure 3I).

Taken together, these analyses demonstrate that NOVA2 contributes to generate AS diversity within specific neuronal types and helps to define specific AS pattern on the same transcripts in different cell types.

### Purkinje cell specific *Nova2* deficiency leads progressive motor discoordination and cerebellar atrophy

We used functional annotation clustering to determine gene ontology (GO) terms associated with the NOVA2-mediated PC-type AS patterns (Figure 3H), identifying synapse related terms (synapse, presynaptic membrane, cell junction, neuronal cell body), intracellular signal transduction, and kinase activity (Figure S4A). This prompted investigation of NOVA2 function in cerebellar PC synaptic and cellular function. When comparing the morphology of control and Nova2-cKO^Pcp2-Cre^ PCs by immunofluorescent staining with appropriate markers, we identified a marked defect in PC dendritic morphology in the absence of *Nova2* (Figure 4A). Nova2-cKO^Pcp2-Cre^ mice also showed noticeable cerebellar atrophy (Figure 4B), motor coordination defects (Figure 4C and Movie S1), PC degeneration (Figure 4D and S4B), loss of synaptic layer thickness accompanied by neuritic swelling (Figure 4E, S4C, and S4D), and reduced spine density (Figure 4F). These results, ranging from histological to cerebellar motor coordination studies to NOVA2 cTag and cKO data reveal NOVA2 biological functions in a specific neuronal cell-type are essential neuronal morphology, activity and survival.

**Figure 4.**
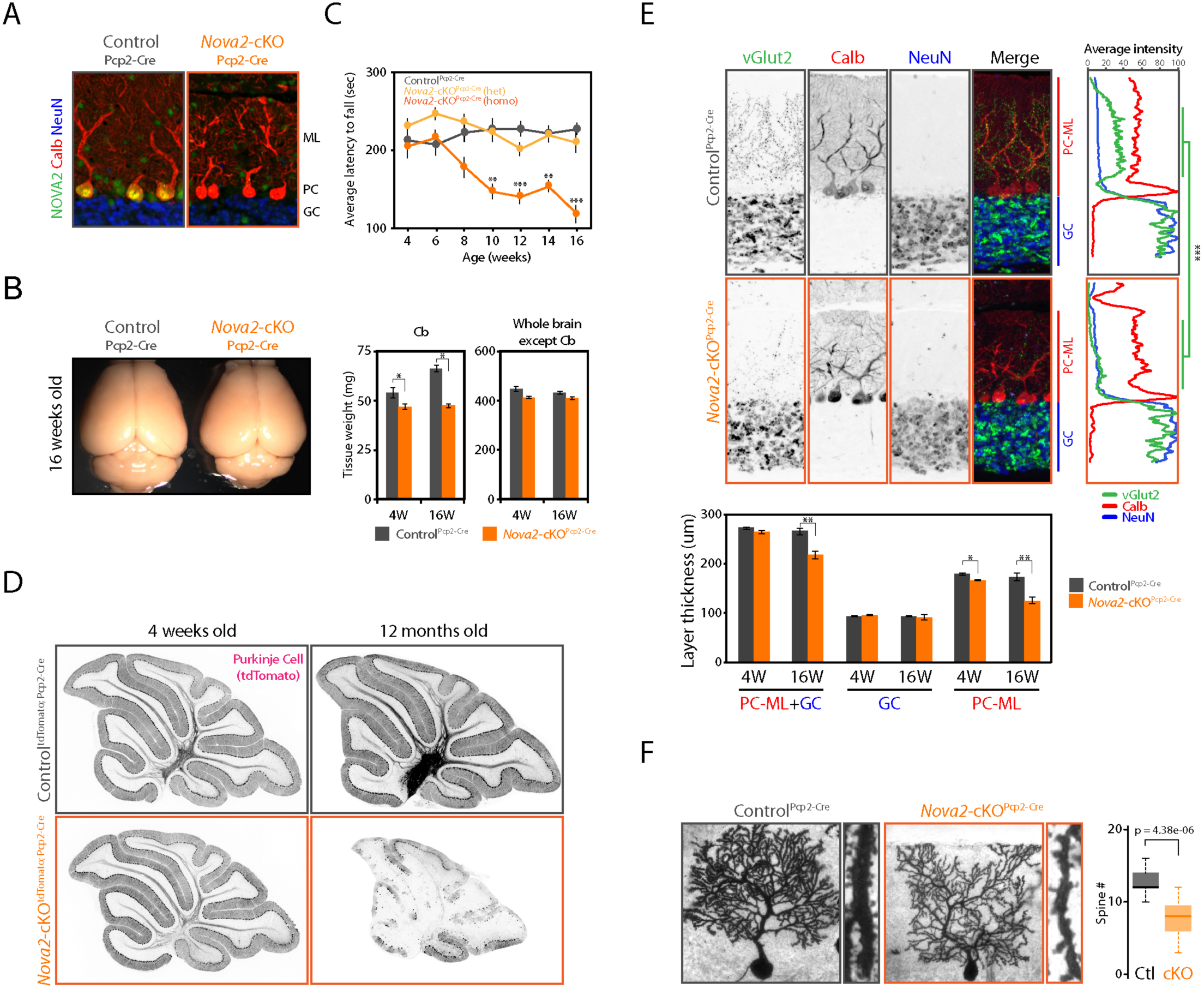
Purkinje cell specific *Nova2* deficiency leads progressive motor discoordination and cerebellar atrophy. (A) Selective NOVA2 depletion from PCs. NOVA2 (*green*), calbindin D28 (calb, *red:* PCs marker), and NeuN (*blue:* GCs marker) immunofluorescent staining images in the 4 weeks old cerebellum of Control^Pcp2-Cre^ and Nova2-cKO^Pcp2-Cre^. (B) Progressive cerebellar atrophy in the PCs specific *Nova2*-cKO mouse. Brain images of 16 weeks old Control^Pcp2-Cre^ and *Nova2*-cKO^Pcp2-Cre^ mouse (*left*). Tissue weight of cerebellum and the other brain regions in 4 weeks old (4W) or 16 weeks old (16W) Control^Pcp2-Cre^ and *Nova2*-cKO^Pcp2-Cre^ mouse (*right*) (n=3, *; p<0.05). (C) Progressive motor coordination defect in the PCs specific *Nova2*-cKO mouse. Rotarod test result at indicated age (weeks) (n>=5, **; p<0.01, ***; p<0.001). (D) PCs-degradation in the 12 months old PCs specific *Nova2*-cKO mouse. ^tdTomato^ immunostaining images with anti-mRFP antibody in 4 weeks old or 12 months old Control^tdTomato; Pcp2-Cre^ and Nova2-cKO^tdTomato; Pcp2-Cre^ mouse. (E) The climbing fiber innervation defect and reduced PCs molecular layer thickness upon PCs specific *Nova2* deficiency. vGlut2 (*green*, climbing fiber terminal marker), calbindin D28 (*red*, PCs marker), and NeuN (*blue*, GCs marker) immunofluorescent staining images in the 16 weeks old Control^Pcp2-Cre^ and *Nova2*-cKO^Pcp2-Cre^ mouse (upper *left*). Quantification analysis of vGlut2, calbindin D28, and NeuN fluorescent intensity in the 16 weeks old Control^Pcp2-Cre^ and Nova2-cKO^Pcp2-Cre^ mouse (n=3, ***; p<0.001) (*upper right*). Quantification analysis of GC and PC-ML layer thickness in 4W or 16W Control^Pcp2-Cre^ and Nova2-cKO^Pcp2-Cre^ mouse (*lower panel*) (n=3, *; p<0.05, **; p<0.01). PC-ML: Purkinje cell and molecular layer. GC: granule cell layer. (F) Reduced PCs spine number upon the NOVA2 depletion. Golgi staining images of 16 weeks old Control^Pcp2-Cre^ and Nova2-cKO^Pcp2-Cre^ mouse (*left*). Quantification of spine numbers per 10 nm dendrite length (neurons number >=14 from three biological replicates) (*right*). “See also Figure S4 and Movie S1.”

### NOVA2 regulates intron retention to serve as a *cis*-acting scaffold element for *transacting* AS factor PTBP2

We showed evidence of NOVA2-mediated AS diversity among the same transcripts expressed in different neuronal types (Figure 2 and Figure 3), leaving open the question of the mechanism by which it does so. To address this, we followed multiple lines of evidence (Coutinho-Mansfield et al., 2007; Markovtsov et al., 2000; Polydorides et al., 2000; Solana et al., 2016; Venables et al., 2013; Zhang et al., 2010; 2016) suggesting functional interactions between RBPs *in vivo*. One such well-studied factor is PTBP2, which plays non-redundant functions with it close paralogue PTBP1 (Vuong et al., 2016) shows differential binding to the corepressor protein Raver1 (Keppetipola et al., 2016), and which has functional interacts with NOVA2 *in vitro* (Polydorides et al., 2000). In fact, we found here that PTBP2 was the *trans*-acting factor whose target AS events (Figure S1C) showed the greatest intersection with NOVA2 target AS events (Figure 5A).

**Figure 5.**
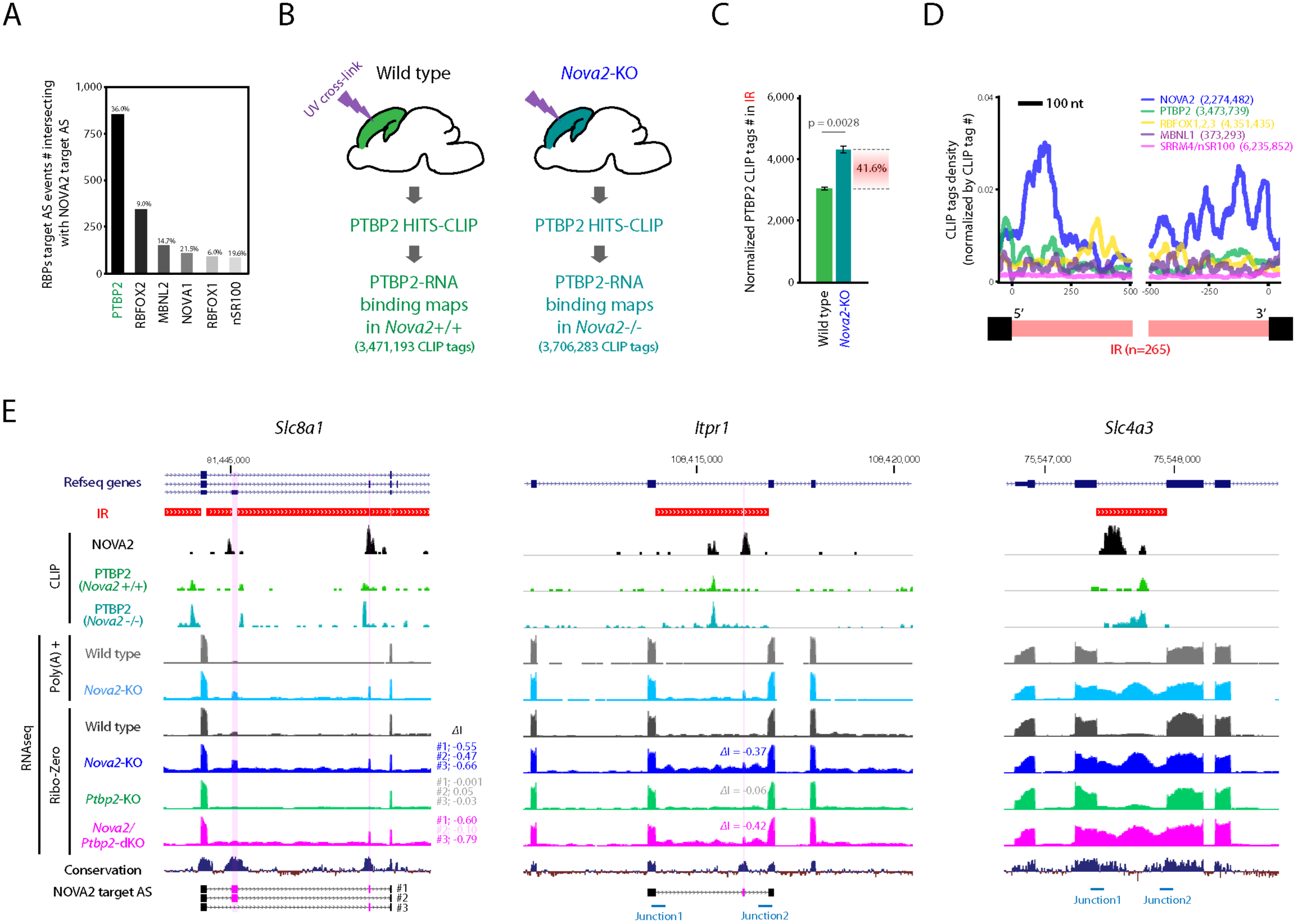
NOVA2 regulates IR to serve as a *cis*-acting scaffold element for AS factor PTBP2. (A) Intersection of NOVA2 and PTBP2 target AS events. The number of RBPs target AS events intersecting with NOVA2 target AS events. (B) Schematics illustrating the overview for PTBP2 CLIP in the presence or absence of NOVA2 *in vivo*. (C) IR recruits trans-acting AS regulator PTBP2. PTBP2 CLIP tag number counted on IR which were normalized by total PTBP2 CLIP tag number of each replicate (n=3). (D) NOVA2 CLIP tags enriched in both 5’ and 3’end of retained introns. (E) UCSC genome browser views (mm10) of examples for PTBP2 CLIP peak changes coupling with IR and AS changes. *Magenta shadow* represents a NOVA2 and/or PTBP2 target AS exon. PTBP2 CLIP height scale were fixed in same scale between wild type and *Nova2*-KO. “See also Figure S5 and Table S1.”

Therefore, we studied PTBP2 dynamic changes in presence or absence of NOVA2 by performing PTBP2 CLIP in the e18.5 cortex of wild type and *Nova2*-KO mice (n=3) (Figure 5B). A total 3,471,193 and 3,706,283 PTBP2 CLIP unique tags were obtained from wild type and *Nova2*-KO cortices, respectively, with comparable CLIP tag genomic distributions (Figure 5B and S5A). Interestingly, we observed that PTBP2 CLIP peak heights in *Nova2*-KO were significantly increased in some neuronal cell-type specific NOVA2 AS targets (e.g. *Itpr1* and *Dab1* in Figure 2H) and were associated with retained intron, leading us to undertake a transcriptome-wide search of NOVA2-dependent intron retention (IR). This identified a total of 265 retained introns in *Nova2*-KO RNA-seq libraries (Figure S5B). These were associated with increased 5’ and 3’ exon-intron junction RNA-seq reads in both Ribo-Zero and poly(A)-selected RNAs, and further confirmed by qRT-PCR interrogating exon-intron junction reads in wild-type and *Nova2*-KO (Figure S5C). Thus these results demonstrate that NOVA2 IR regulation occurred in *cis* in poly(A) mRNAs. PTBP2 CLIP tag numbers in retained introns were significantly increased in *Nova2*-KO (Figure 5C) but were comparable in other introns or elsewhere in the same transcripts (Figure S5D).

Interestingly, in wild-type mouse brain, NOVA2 CLIP tags and peaks were enriched on the 5’ and 3’ end of NOVA2-dependent retained introns but not in the other introns within the same transcripts (Figure 5D and S5E). The enrichment of NOVA2 was specific, as we found no CLIP tag enrichment of several other RBPs, including PTBP2, RBFOX1/2/3, MBNL1, and nSR100 in these 265 introns (Figure 5D). Taken together, the results indicate that NOVA2 is specifically associated with IR removal, likely in conjunction with its binding to local 5’ and 3’ introns. Overall, these results demonstrate that NOVA2 regulated IR induces dynamic PTBP2 binding alternation in it.

To determine whether increased PTBP2 binding to retained introns had a functional impact on AS, we analyzed RNA-seq from e18.5 cortex of *Nova2/Ptbp2* double-KO (dKO) mice. We found total 3,415 AS events as significant AS events changing in *Nova2/Ptbp2*-dKO (FDR<0.05, |*Δ*I|>=0.1) relative to wild-type mice and 876 of 3,415 AS events were specific AS events changed in *Nova2/Ptbp2*-dKO (Figure S5F). We observed two kinds of instances of AS events which were additively or synergistically regulated by both NOVA2 and PTBP2 (Figure S5G), indicating that NOVA2 and PTBP2 can co-regulate the same AS events. In AS regions that overlapped NOVA2-suppressed IR, PTBP2 CLIP peak height was consistently increased in *Nova2*-KO (Figure 5E and S5H). AS changes in *Nova2*-KO were attenuated in *Slc8a1^#2^, Mcf2l*, and *Lrrfip1* and enhanced in *Itpr1, Agrn, Ppp3cb*, and *Bin1* when PTBP2 was depleted from *Nova2*-KO brain (in *Nova2/Ptbp2*-dKO) (Figure 5E, S5G, and S5H). These results suggest a role for IR as a scaffold for PTBP2 binding with consequences on AS regulation, whereby PTBP2 binding to NOVA2-suppressed IR is increased in the absence of NOVA2 *in vivo*. Taken together, these results demonstrate that the *trans*-acting AS factor NOVA2 regulates IR as a *cis*-acting scaffolding platform for the other *trans*-acting AS factor.

### Multiple NOVA2-mediated target RNA metabolism coupling to IR

We found that NOVA2-dependent IR was unique for each individual neuronal type and coupling to AS change, as seen with *Itpr1* and *Slc8a1* (Figure 6A). NOVA2-dependent *Slc8a1*^#^*^1^*, *Slc8a1^#2^* and *Itpr1* AS events were significantly changed only in cortical excitatory neurons but not in cortical and cerebellar inhibitory neurons, which were coupled to selective NOVA2-dependent IR increases in cortical excitatory neurons. The insertion of *Itpr1* NMD exon were embedded between *Itpr1* IR1 and IR2 (Figure 6A). *Itpr1* mRNA was significantly reduced only in PC specific *Nova2*-cKO when compared with controls (FDR<0.01, log2 FC <= -1), coincident with selective NOVA2 cTag-CLIP binding in the *Itrp1* 3’ UTR of PCs (Figure 6A), but not in cortical excitatory and inhibitory neuron transcripts, despite the retention of the NMD exon in the cortical excitatory *Itpr1* transcript (Figure 6A). Interestingly, ITPR1 protein levels assessed by Western blot were reduced in both whole cortical *Nova2*-KO and whole cerebellar *Nova2-*cKO^Pcp2-Cre^ tissues (Figure 6B). These results, summarized in Figure 6C, show the dual function of NOVA2 on *Itpr1* RNA metabolism between different neuronal types, indicating that NOVA2-dependent IR removal contributes to generate neuronal cell-type specific mRNA diversity and RNA metabolism diversity (Figure 6D).

**Figure 6.**
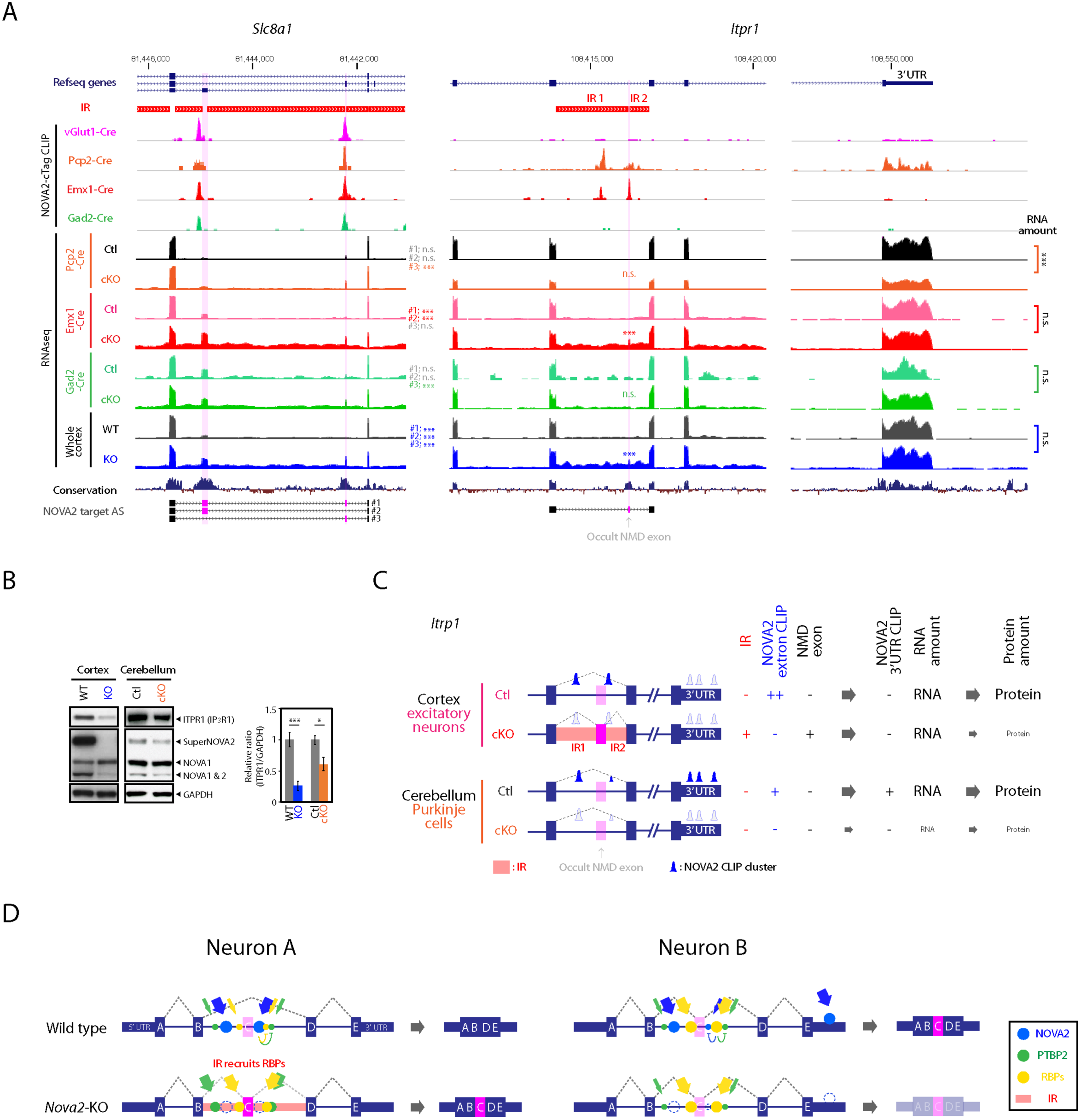
Multiple NOVA2-mediated target RNA metabolism coupling to IR. (A) Neuronal subtype selective and NOVA2-dependent AS changes coupling with IR. UCSC genome browser views (mm10) of examples (*Slc8a1* and *Itpr1*). ***; FDR<0.001. (B) Decreased ITPR1 protein levels both in the cortex and cerebellum. Immunoblot analysis with anti-ITPR1 antibody in e18.5 cortex of wild type and *Nova2*-KO and in 16 weeks old whole cerebellum of Control^Pcp2-Cre^ and Nova2-cKO^Pcp2-Cre^ (n=3, *; p<0.05, ***; p<0.001). (C) Summary of *Itpr1* mRNA metabolism mediated by NOVA2. (D) Model Schematics for neuronal subtype specific AS and its regulatory mechanism. Neuronal subtype selective IR coupling with NOVA2-dependent and neuronal subtype selective AS changes recruit other RBPs, generating AS diversity between different neuronal cell-types.

## DISCUSSION

In the present work, we discover that a single splicing factor mediates AS diversity across different CNS regions, individual neuronal types, and with different mechanisms. These results rely on the establishment of a new high-resolution CLIP platform technology, “cTag-CLIP”, for generating regulatory RBP-RNA interaction maps with neuronal type specific resolution from intact tissue *in vivo*. Bioinformatic integration of neuronal type specific RBP-RNA maps with corresponding RNA-seq datasets provides the first evidence for the diversity of RBP-RNA binding properties and the discovery of neuronal type selective RBP target AS events, and new means of regulating metabolism of the same transcript in different neuronal types.

Analysis focused on cerebellar PCs revealed that NOVA2 characterizes a subset of PCs-type AS patterns and contributes to spawn alternatively spliced transcripts between cerebellar PCs and GCs. RNA processing diversity including AS among different neuronal types might be spawned by the difference of RBPs expression ratio in each neuronal type, of RBPs or target RNA modification, and/or of unknown RBPs co-factor. It is supported by the different expression levels of AS regulator (e.g. NOVA1, NOVA2, RBFOX3, and PTBP2) across the CNS areas and neuronal types (Figure 1C). The diverse expression of the other RBPs (e.g. RBFOX, PTBP, and ELAVL/Hu proteins), which is known as AS and translation regulatory factor, has been also reported (Gehman et al., 2012; Ince-Dunn et al., 2012; Makeyev et al., 2007). Future *in vivo* cell-type specific approach would provide the further evidences of RBPs-mediated RNA regulation diversity, including not only AS and RNA stability but also translation and RNA modification, across different neuronal types. These *in vivo* cell-type specific analysis providing the unique RNA metabolism information for individual cell populations would contribute to understand diseases mechanism, such as amyotrophic lateral sclerosis (ALS), frontotemporal dementia (FTD), spinocerebellar ataxia (SCA), and spinal muscular atrophy (SMA), which are strongly correlating with abnormality of RBP and/or RNA metabolism. Applying our *in vivo* cTag strategy into vulnerable cell-type to disease may answer the questions why there is vulnerable cell-type to specific disease and how functions of these specific cell-types are disorganized in disease. Overall, our *in vivo* cell-type specific approach offers novel insights into cell-specific RNA biology within intact living tissues.

In the past few years, low translation efficiency of transcript containing retained introns has been reported (Braunschweig et al., 2014; Gill et al., 2017; Pendleton et al., 2017). We discover that NOVA2 acts as a *cis*-acting AS element to regulate intron retention, which subsequently serve as cell-specific scaffolds with which to alter RBP-RNA dynamic interactions of another cell-type specific *trans*-acting splicing factor, PTBP2. This result provides novel insights into the regulation and biological function of intron retention.

## AUTHOR CONTRIBUTIONS

Y.S. and R.B.D. wrote the manuscript and designed experiments. Y.Y. generated *Nova2*-cKO mice. I.Z.S performed rotarod test. J.J.F. prepared all RNA-seq libraries. Y.T performed and helped RT-PCR experiments. D.D.L. and Y.S. performed PTBP2 CLIP. Y.S. generated *Nova2-* cTag mice and performed all other experiments and data analysis.

## ACKNOWLEDGEMENTS

We thank R.B. Darnell lab members for suggestions, expertise, and feedback. R.B.D. is an Investigator of the Howard Hughes Medical Institute. Support was provided by the National Institutes of Health (NS034389, NS081706, NS097404, and 1UM1HG008901) and the Simons Foundation (SFARI 240432) to R.B.D.. Y.S. is a recipient of JSPS postdoctoral fellowship for research abroad and Naito Foundation.

## STAR METHODS

### KEY RESOURCES TABLE

Will be uploaded separately.

### LEAD CONTACT FOR REAGENT AND RESOURCE SHARING

Further information and requests for resources and reagents generated in this study should be directed to and will be fulfilled by the Lead Contact, Robert B. Darnell.

### EXPERIMENTAL MODEL AND SUBJECT DETAILS

#### Generation of *Nova2-cTag* and *Nova2*-cKO mice lines

Nova2-cTag and *Nova2*-cKO targeting vectors were generated through standard restriction cloning. Nova2-cTag targeting vector contains the endogenous *Nova2* exon 4 preserving 6.7 kb 3’ UTR, triple synthetic poly(A) sites preceding *FRT-NEO-FRT* cassette, and last coding exon of *Nova2* fused to AcGFP sequence (Figure S2A). *Nova2*-cKO targeting vector contains *Nova2* exon1 preceding *FRT-NEO-FRT* cassette (Figure S2F). Each construct was electroporated into Bruce 4 ES cells. Correctly targeted ES cells were injected into C57BL/6J blastocysts to screen for chimeras. Chimeric males were bred to C57BL/6J females to generate either heterozygous Nova2-cTag or *Nova2*-cKO. The *FRT-NEO-FRT* cassette was removed by breeding heterozygous *Nova2*-cTag or *Nova2*-cKO to ACTB-FLPe mice.

#### Animal experiments

All procedures were conducted according to the Institutional Animal Care and Use Committee (IACUC) guidelines at the Rockefeller University. C57BL/6J (Stock No. 000664), ACTB-FLPe (Stock No. 005703), ^tdTomato^ reporter (Stock No. 007914), CMV-Cre (Stock No. 006054), Emx1-Cre (Stock No. 005628), Gad2-Cre (Stock No. 010802), vGlut1-Cre (Stock No. 023527), and Pcp2-Cre (Stock No. 004146) mice were obtained from Jackson Lab. *Nova2*-KO and *Ptbp2-* KO mice were previously described (Licatalosi et al., 2012; Ruggiu et al., 2009; Saito et al., 2016; Yano et al., 2010). Mice were housed up to 5 mice per cage in a 12 hour light/dark cycle. For genotyping PCR experiment, mouse genomic DNA was extracted from a tail. The primer sequences are listed in Table S2. Mice of both sexes were pooled for the study. Sex-specific effects were not tested. 4–16 weeks old *Nova2*-cKO^wt/wt^; Pcp2-Cre^+/-^ (Control^Pcp2-Cre^), *Nova2-*cKO^Flox/wt^; Pcp2-Cre^+/-^ (*Nova2*-cKO^Pcp2-Cre^ het), and *Nova2*-cKO^Flox/Flox^; Pcp2-Cre^+/-^ (*Nova2-* cKO^Pcp2-Cre^ homo) mice on a C57BL/6 background were used for rotarod test.

For fluorescence activated cell sorting (FACS), age-matched e18.5 mice cortex of Control^tdTomato; Emx1-Cre^, Control^tdTomato; Gad2-Cre^, *Nova2*-cKO^tdTomato; Emx1-Cre^, and *Nova2*-cKO^tdTomato; Gad2-Cre^, were digested with 2 mg/mL papain (Worthington; PAPL LSS03119)/Hibernate E minus calcium solution (BrainBits) (n=3). The resulted single cell suspension were subjected to FACS to collect ^tdTomato^(+) and (-) cells.

The tissue region, age, genotype, and number used for CLIP experiment were listed in Table S3. Manually dissected tissue was homogenized 2–3 times by using 1 mL syringe with 20G ×1 ½ needle in 1 mL ice-cold PBS. The tissue block solution was transferred to 6 well dishes (FALCON; 353046) on ice for UV-crosslinking. The UV-crosslinked tissue block was spun at 4°C and the pellet was stored at -80°C until usage (three biological replicates for each CLIP).

For immunohistochemistry and Golgi staining, e18.5, 3 weeks, 4 weeks, 16 weeks, or 1 year old mice were perfused with PBS and fixed with 4% praraformaldehyde (PFA)/PBS at 4 degrees overnight, and sequentially replaced to 15% sucrose/PBS and 30% sucrose/PBS at 4 degrees for cryo-protection, then embedded in OCT compound and stored at -80°C until usage.

For manual dissection of cerebellar Purkinje cell layer, cerebellar sections (200 μm) were prepared from 4 weeks old cerebellum of Control^tdTomato; Pcp2-Cre^ and Nova2-cKO^tdTomato; Pcp2-Cre^ by using tissue chopper (McILWAIN TISSUE CHOPPER) and were pooled in ice cold Hebernate EB complete (BrainBits). Purkinje cell layer was manually dissected from the sections by using tungsten dissection needle (ROBOZ; RS-6065) under the SteREO Discovery.V12 fluorescent microscope (Zeiss), snap-frozen in liquid nitrogen, and stored -80°C until usage.

### METHOD DETAILS

#### Antibodies

Primary antibodies used for CLIP, immunofluorescent staining (IF), and western blots (WB) were: mouse anti-GFP ((Heiman et al., 2014), clones 19F7 and 19C8, cTag-CLIP), rat anti-GFP (nacalai tesque, GF090R, IF: 1/1,000), mouse anti-GFP (Santa Cruz, sc-9996, WB: 1/1,1000), rabbit anti-mRFP/^tdTomato^ (ROCKLAND, 600–401–379, IF: 1/2,000), rabbit anti-NeuN/RBFOX3 (Millipore, ABN78, IF: 1/1,000), guinea pig anti-NeuN/RBFOX3 (Millipore, ABN90P, IF: 1/1,000), goat anti-NOVA2 (Santa Cruz, sc-10546, IF; 1/500, WB; 1/2,000, CLIP), rabbit anti-NOVA1 [EPR13847] (abcam, ab183024, IF: 1/1,1000), human anti-pan NOVA (anti-NOVA paraneoplastic human serum, WB: 1/5,000), rabbit anti-PTBP2 (Polydorides et al., 2000, IF:1/5,000, CLIP), guinea pig anti-vGlut1 (Synaptic System, 135 304, IF:1/2,000), rabbit anti-vGlut2 (Synaptic System, 135 403, IF:1/2,000), mouse anti-VGAT (Synaptic System, 131 011, IF: 1/1,000), rabbit anti-Calbindin D28 (Millipore, AB1778, IF: 1/1,000), goat anti-Calbindin D28 (Santa Cruz, sc-7691, IF: 1/500), rabbit anti-ITPR1/InsP3R, type1 (Millipore, ABS55, IF:1/2,000), mouse anti-GAPDH (Santa Cruz, sc-32233, WB: 1/1,000).

#### Fluorescence activated cell sorting

Age-matched e18.5 mice cortex of Control^tdTomato; Emx1-Cre^, Control^tdTomato; Gad2-Cre^, *Nova2-* cKO^tdTomato; Emx1-Cre^, and *Nova2*-cKO^tdTomato; Gad2-Cre^ were digested with 2 mg/mL papain (Worthington; PAPL LSS03119) in Hibernate E minus calcium solution (BrainBits) for 15 min at 37°C (n=3 for each genotype), added 1/5 volume of 1 mg/mL DNase I/Hibernate E minus calcium solution, incubated for 5 min at 37°C, and transferred on ice. After 2 times wash with ice cold Hibernate EB solution (BrainBits), tissues were gently pipetted 7–10 times with 1,000 μL tip. Large clumps of cells were removed using a cell strainer (BD Biosciences). The single cell suspensions were centrifuged for 2 min at 4°C and re-suspended with ice cold 3% FBS/PBS with 100 ng/mL DAPI. Large clumps of cells were removed using a cell strainer (BD Biosciences) again. ^tdTomato^ positive cells or negative cells were sorted into each tube filled with Trizol LS (ambion; 10296010) on a BD FACSAria Cell Sorter (BD Biosciences) and RNA was isolated for analysis. DAPI positive cells were excluded from samples.

### RT-PCR

RNA was reverse transcribed using SuperScript III First-Strand synthesis System for RT-PCR (Invitrogen; 18080–051). qRT-PCR was performed with FastStart SYBR Green Master (Roche; 04 673 492 001) on BIO-RAD Touch™ Real-Time PCR Detection System. The following programme was carried to 40 cycles: 30 s 95 °C (denaturation); 30 s 58 °C (annealing); and 20 s 72 °C (extension). Results were analysed by ΔΔCt, using *Actb* mRNA for normalization. Semi-quantitative RT-PCR was performed as previously described (Saito et al., 2016).

#### SDS-PAGE and Western blots

E18.5 whole cortex or P28 whole cerebellum were lysed in PXL buffer (1x PBS; 0.1% SDS; 0.5% Na-DOC; 0.5% NP-40) and lysates (10 ug protein/lane) were separated by NuPAGE™ 4–12% Bis-Tris Protein Gel (Invitrogen; NP0323BOX) and transferred to nitrocellulose blotting membrane (GE Healthcare Life science; 10600002). Antibodies used were listed above. Western blots were quantified normalizing each lane by the GAPDH signal to control for loading differences.

#### Immunohistochemistry and Golgi staining

Immunohistochemistry was performed as previously described (Saito et al., 2016). Briefly, 4 weeks old mice brains were perfused with PBS and fixed with 4% praraformaldehyde (PFA)/PBS at 4 degrees overnight or e18.5 mice brain were fixed with 4% PFA/PBS at 4 degrees overnight, and sequentially replaced to 15% sucrose/PBS and 30% sucrose/PBS at 4 degrees for cryo-protection, then embedded in OCT compound. Frozen brains were sliced into 30 μm (for 4 weeks old mice) or 80 μm (for e18.5 mice) thick sections on a cryostat (CM3050S, LEICA). Brain sections were subjected to immunohistochemistry. These samples were washed three times with PBS at room temperature (RT), incubated with 0.2% Triton X-100/PBS for 15 min at RT, blocked with 1.5% normal donkey serum (NDS)/PBS for 1 hr at RT, and then incubated overnight at 4 degrees with primary antibodies in 1.5% NDS/PBS followed by incubation with Alexa 488, 555 or 647 conjugated donkey secondary antibodies (1:1000) in 1.5% NDS/PBS. Golgi staining were performed according to FD Rapid GolgiStain™ Kit (FD NeuroTechnologies, INC.) manual. Images of specimens were collected by BZ-X700 (KEYENCE) microscope.

#### Rotarod test

Mice (4, 6, 8, 10, 12, 14, and 16 weeks of age, n>=5 per group) were allowed to acclimate in a behavioral room for 1 hr and then placed on an accelerating rotarod (Med-Associates). On day one only mice were trained. Mice were considered trained when they could stay on the apparatus rotating at 4 RPM without falling for 1 min or until they fell off 5 times. During test trails animals were placed on apparatus rotating at 4 RPM, which was gradually increased to 40 RPM over 5 mins. Latency to fall (in seconds) from the rotating bar was recorded. Mice were rested for 5 min and repeated two additional times per day for three consecutive days. A mouse’s latency to fall for each day was recorded as the mean latency of the three consecutive trials and data are reported ± SEM.

#### RNA-seq library preparation and analysis

mRNA-seq libraries were prepared from Trizol extracted RNA following Illumina TruSeq protocols for Ribo-zero or poly-A selection, fragmentation, and adaptor ligation. Multiplexed libraries were sequenced as 125 nt paired-end runs on HiSeq-2500 platforms at New York Genome Center. These raw datasets and processed data files have been deposited into Gene Expression Omnibus (GSE103316). The publically available RNA-seq and TRAP datasets used in this study were listed in Table S3. Reads were aligned to the mm10 build of the mouse genome using OLego (Wu et al., 2013), and AS and gene expression were analyzed using the Quantas pipeline (Charizanis et al., 2012; Yan et al., 2015). Transcripts isoform abundance in RNA-seq datasets was estimated using Kallisto (Bray et al., 2016) with customized aggregation index. The mRNA abundance change was assessed by differential analysis of raw sequencing counts in edgeR using TMM methodology. For IR analysis, exon and intron information was obtained from Refseq and Quantas AS events database, and removed duplicated exons and introns. RNA-seq reads mapped on only either intron or exon-intron junction were used for counting the number of reads on either intron or exon-intron junction, respectively. Counted intron or exon-intron junction reads were normalized with total reads counted on its gene, including UTRs, CDSs, and introns, for each biological replicate, and subjected to differential analysis (edgeR). Short introns (less than 250 nt) and introns of low expression genes (less than 100 RNA-seq reads counted on gene for each replicate were excluded from analysis. 265 introns (intron reads; log_2_FC(*Nova2*-KO/wild-type)>0.6 and p<0.01 in both Ribo-zero and Poly(A)+ libraries, and 3’ and 5’ exon-intron junction reads; log_2_FC(*Nova2*-KO/wild-type)>0.6 and p<0.01 in either Ribo-zero or Poly(A)+ library) were used as NOVA2-dependent IR (GSE103316). Low coverage introns (average exon-intron junction reads less than 5 reads in *Nova2*-KO) were excluded from analysis.

#### Multiplexed CLIP library preparation

PTBP2 CLIP in the e18.5 cortex of wild type and *Nova2*-KO and NOVA2 CLIP in P28 cerebellum were performed generally following previous work (Licatalosi et al., 2012; Saito et al., 2016) combining with multiplexed step (Luna et al., 2015). The publically available CLIP datasets used in this study were listed in Table S3 (NOVA2, MBNL1, RBFOX1/2/3, and nSR100/SRRM4 CLIP). NOVA2 cTag-CLIP was performed as previously described (Ule et al., 2005a) with the following modifications. UV-crosslinked NOVA2-AcGFP::RNA complex extracted using ice cold PXL buffer were immunoprecipitated using mouse monoclonal anti-GFP clones 19F7 (12.5 μg) and 19C8 (12.5 μg) mixture with 200 μL of Dynabeads™ Protein G magnetic beads (Thermo Fisher SCIENTIFIC; 10009D). The mouse brain samples used for NOVA2 cTag-CLIP was listed in Table S3. For controls, mouse brain (no UV irradiation, *Nova2*-cTag^Cre(-)^, and overdigested control) were used. Prior to phosphatase treatment, beads were washed 2 times with each in a series of wash buffers (see below). NOVA2-AcGFP bound RNA fragments were ligated to a degenerate 5’ RNA linker. The two-step PCR amplification was performed with AccuPrime™ Pfx Polymerase (Invitrogen). cDNA was then amplified with MSFP3 and indexed MSFP5 primers pair (Luna et al., 2015). Samples were sequenced (single-end 75 nt) on the MiSeq system (Illumina).

Wash buffers; Wash buffer 1 (PXL) [1xPBS; 0.1%SDS; 0.5%Na-DOC; 0.5% NP-40], Wash buffer 2 [15 mM Tris, pH7.4; 5 mM EDTA; 2.5 mM EGTA; 1% Triton X-100; 1% Na-DOC; 0.1% SDS; 120 mM NaCl; 25 mM KCl], Wash buffer 3 [15 mM Tris, pH7.4; 5 mM EDTA; 2.5 mM EGTA; 1% Triton X-100; 1% Na-DOC; 0.1% SDS; 1 M NaCl], Wash buffer 4 [15 mM Tris, pH7.4; 5 mM EDTA], Wash buffer 5 [50 mM Tris, pH7.4; 10 mM MgCl_2_; 0.5% NP-40].

#### Analysis of CLIP data

Analysis of CLIP data were carried out similar to previous reports (Moore et al., 2014; Shah et al., 2017). Data was visualized with UCSC genome browser (http://genome.ucsc.edu/). To reduce mis-alignments due to sequencing errors, reads were initially filtered based on quality score (≥20 in the degenerate linker region; average of ≥20 in the remaining read). Exact sequences were collapsed to remove PCR duplicates and de-multiplexed. The degenerate barcode was removed and the 3’ linker was trimmed. CLIP reads were mapped on the mm10 build of the mouse genome by novoalign (www.novocraft.com). Only unique tags were used for subsequent analyses. All scripts used in the analysis including the peak finding algorithm and more information can be publically obtained at http://zhanglab.c2b2.columbia.edu/index.php/Resources.

### QUANTIFICATION AND STATISTICAL ANALYSIS

#### Alternative splicing quantification

RNA-seq data were aligned with OLego de-novo splicing aligner. AS were quantified using Quantas. Quantas calculates exon inclusion rate and differential exon inclusion rate between two groups (*Δ*I). Fisher exact test in Quantas was used to call FDR adjusted P values. We used FDR < 0.05 and *Δ*I > |0.1| as significant AS events.

### DATA AND SOFTWARE AVAILABILITY

The GEO accession number for NOVA2 cTag-CLIP and RNA-seq originated from this study is GSE103316. Supplemental Figures, Tables, and Movie Legends

### Supplemental Figures, Tables, and Movie Legends

**Figure S1.**
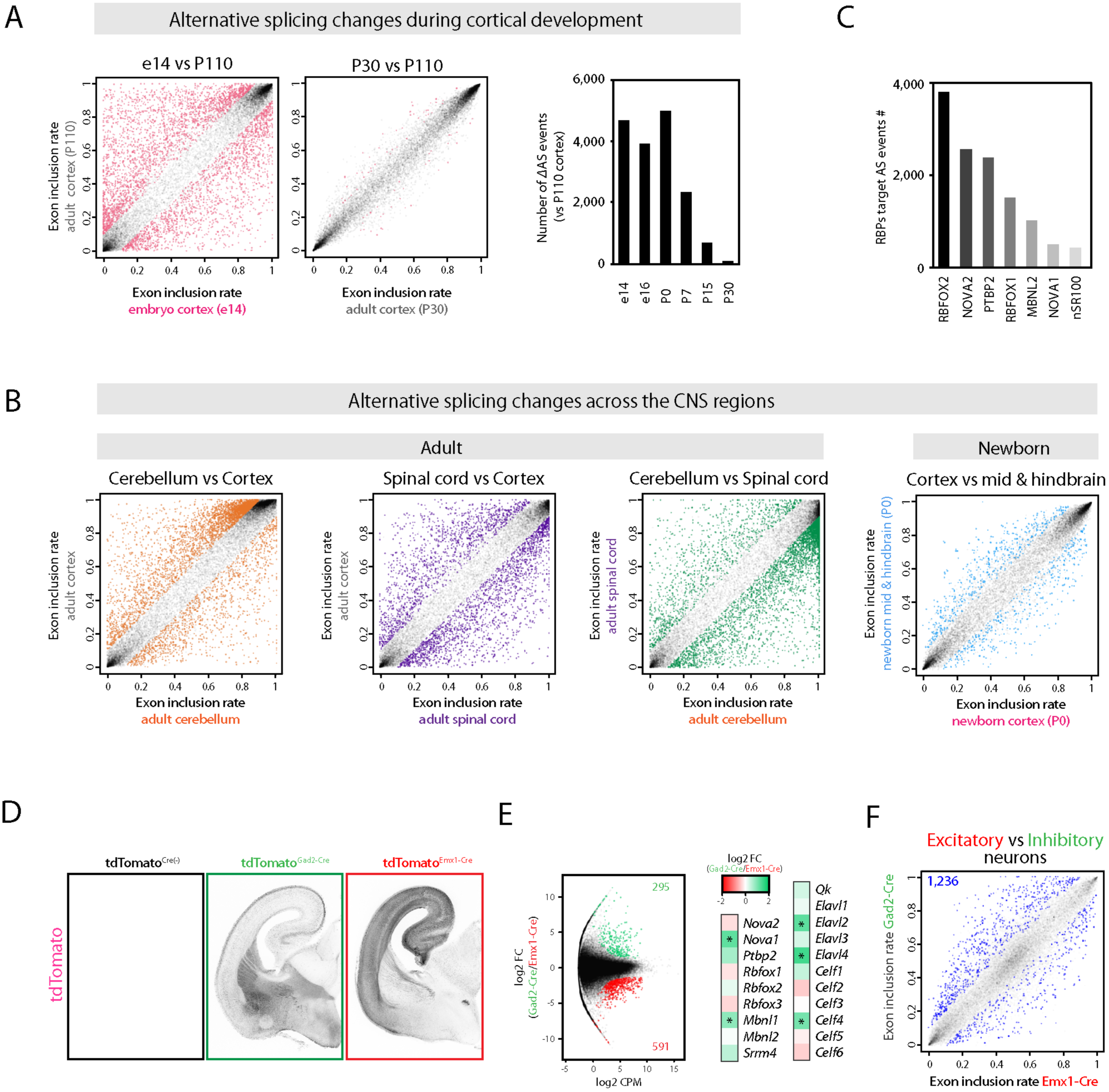
Related to Figure 1. Diversity of AS across the CNS regions and neuronal types. (A) AS changes during mouse cortical development. The examples of developmentally regulated cassette-type AS events in the mouse cortex between e14 or P30 and P110 mouse cortex. *Pink* dot indicates the significantly changed AS events (FDR<0.05, |*Δ*I|>=0.1) (*Left* two scatter plot). Summary of total cassette-type AS number that significantly changed when compared with P110 mouse cortex (FDR<0.05, |*Δ*I|>=0.1) (*right*). (B) AS diversity across the CNS regions. Scatter plots show the exon inclusion rate of cassette-type AS events between the CNS regions. Each *Orange, purple, green*, or *light blue* dot indicates the significantly changed AS event between adult cortex and cerebellum, adult cortex and spinal cord, adult cerebellum and spinal cord, and newborn cortex and the mixture of midbrain, hindbrain, and cerebellum, respectively (FDR<0.05, |*Δ*I|>=0.1). (C) Total AS events number significantly changing in each RBP-KO mouse brain. AS changes were identified from publically available RNA-seq datasets (FDR<0.05, |*Δ*I|>=0.1). (D) Cre dependent ^tdTomato^ expression. ^tdTomato^ immunofluorescence staining images in e18.5 brain sections of Control^tdTomato; Cre(-)^, Control^tdTomato; Gad2-Cre^, and Control^tdTomato; Emx1-Cre^ mouse. (E) RBPs mRNA expression difference between neuronal linages. Asterisks show the significant changes (FDR<0.05, |log2FC|>=1). (F) AS comparison between e18.5 Gad2-Cre and Emx1-Cre linage. *Blue* dot indicates a significantly changed AS event determined by RNA-seq (FDR<0.05, |*Δ*I|>=0.1).

**Figure S2.**
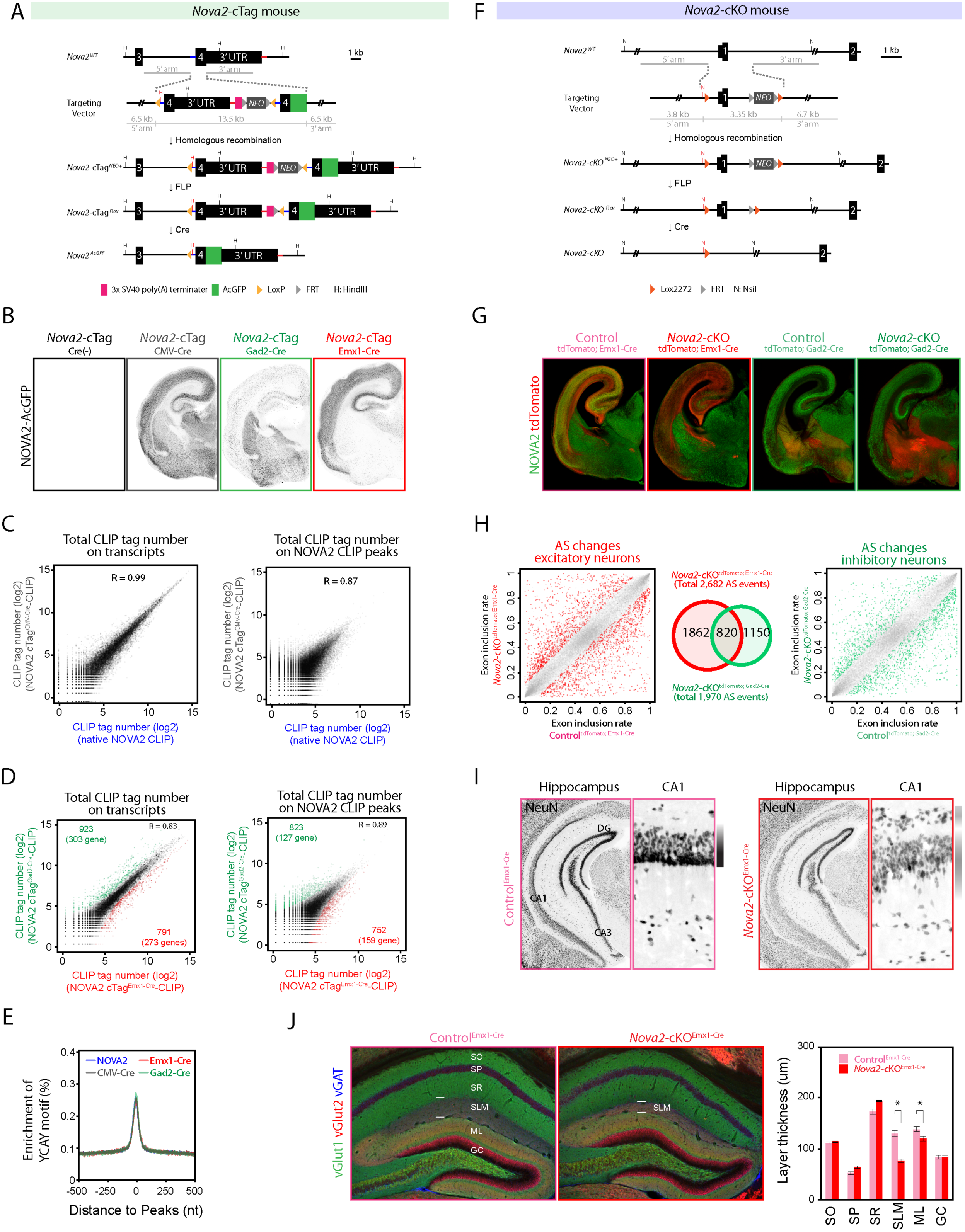
Related to Figure 2. Mice models development providing neuronal linage selective and transcriptome-wide NOVA2 analysis. (A) Targeting strategy for generating the knock-in Nova2-cTag mouse. FLP: Flippase. FRT: Flippase recognition target. NEO: Neomycin resistance gene. (B) Cre-dependent NOVA2-AcGFP protein expression. NOVA2-AcGFP immunofluorescent staining images with anti-GFP antibody in e18.5 Nova2-cTag^Cre(-)^, *Nova2-* cTag^CMV-Cre^, Nova2-cTag^Gad2-Cre^, and Nova2-cTag^Emx1-Cre^ mice brain. (C) The comparison of NOVA2 and NOVA2-AcGFP binding property to RNA. Scatter plots show the correlation of read counts between conventional NOVA2-CLIP to NOVA2 cTag^CMV-Cre^-CLIP. Correlation of total CLIP read counts on transcripts (*Left*) and on NOVA2 CLIP peaks (*right*). Each dot represents a transcript (*left*) or CLIP peak (*right*). R: correlation coefficient. (D) The comparison of neuronal linage specific NOVA2 cTag-CLIP between cortical excitatory (Emx1-Cre) and inhibitory neurons (Gad2-Cre). Scatter plots show the correlation of read counts between NOVA2 cTag^Gad2-Cre^-CLIP to NOVA2 cTag^Emx1-Cre^-CLIP. Correlation of total CLIP read counts on transcripts (*Left*). Correlation of total CLIP read counts on NOVA2 CLIP peaks (*right*). Each *black* dot represents a comparable transcript (*left*) or CLIP peak (*right*) between two neuronal linages. Each *green* or *red* dot represents a significantly enriched transcript (*left*) or CLIP peak (*right*) in NOVA2 cTag^Gad2-Cre^-CLIP or NOVA2 cTag^Emx1-Cre^-CLIP, respectively (FDR<0.05, |log2 FC|>=1). R: correlation coefficient. (E) Enrichment of YCAY motif near conventional NOVA2 CLIP and NOVA cTag-CLIP peaks. (F) Targeting strategy for generating the *Nova2-* cKO mouse. FLP: Flippase. FRT: Flippase recognition target. *NEO:* Neomycin resistance gene. (G) NOVA2 depletion from selective neuronal populations. NOVA2 (*green*) and ^tdTomato^ (*red*) immunofluorescent staining images in the e18.5 Control^tdTomato; Emx1-Cre^, Nova2-cKO^tdTomato; Emx1-Cre^, Control^tdTomato; Gad2^-^Cre^, and Nova2-cKO^tdTomato; Gad2-Cre^ mice brain. (H) NOVA2 mediated AS events in restricted neuronal population. *Left and right* scatter plots show the exon inclusion rate of the e18.5 cortical Control^tdTomato; Emx1-Cre^, Nova2-cKO^tdTomato; Emx1-Cre^, Control^tdTomato; Gad2-Cre^, and *Nova2*-cKO^tdTomato; Gad2-Cre^. Each *grey* dot represents an AS event. Each *Red* and *green* dot indicates the significantly changed AS event in Emx1-Cre or Gad2-Cre lineage, respectively (FDR<0.05, |*Δ*I|>=0.1). *Middle* Venn diagram representing NOVA2 target AS events number overlapping or differentially regulated between two neuronal-linages. (I) The disorganization of hippocampal laminar structure in Nova2-cKO^Emx1-Cre^ mice. NeuN immunostaining in the hippocampus of 3 weeks old Control^Emx1-Cre^ (*left, pink*) or Nova2-cKO^Emx1-Cre^ mice (*right, red*). (J) CA1 molecular layer disorganization in 3 weeks old Emx1-Cre; *Nova2*-cKO mice. Hippocampal layer visualized by immunostaining with anti-vGlut1, vGlut2, and vGAT antibodies in 3 weeks old Control^Emx1-Cre^ or *Nova2*-cKO^Emx1-Cre^ mice (*left images*). Quantification and comparison of hippocampal layer thickness between Control^Emx1-Cre^ or *Nova2*-cKO^Emx1-Cre^ mice (*right*). *: p<0.05. SO; stratum oriens, SP; apical dendrites stratum pyramidal, SR; stratum radiatum, SLM; stratum lacunosum moleculare, ML; molecular layer, GC; granule cell.

**Figure S3.**
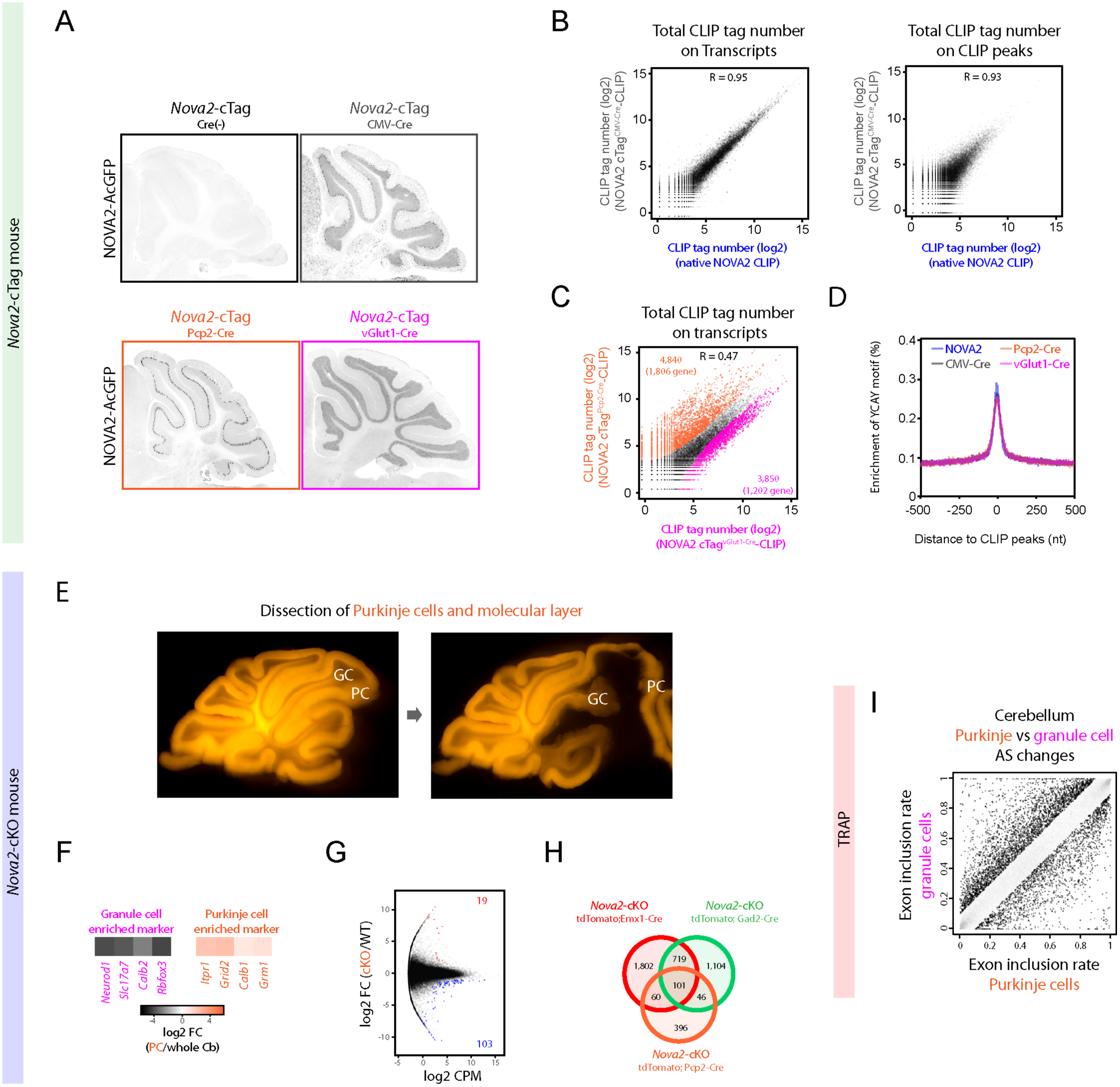
Related to Figure 3. Neuronal type specific analysis in adult cerebellum. (A) Cre-dependent NOVA2-AcGFP expression in cerebellum. Mouse brain slices prepared from 4 weeks old Nova2-cTag mice mated with CMV-Cre, Pcp2-Cre (Purkinje cell), vGlut1-Cre (granule cell) driver mice were subjected to immunostaining with anti-GFP antibody. (B) The comparison of NOVA2 and NOVA2-AcGFP binding property to RNA in adult cerebellum. Scatter plots show the correlation of read counts between conventional NOVA2-CLIP to NOVA2 cTag-CLIP (CMV-Cre). Correlation of total CLIP read counts on transcripts (*Left*) and on NOVA2 CLIP peaks (*right*). Each dot represents a transcript (*left*) or CLIP peak (*right*). R: correlation coefficient. (C) The comparison of neuronal type specific NOVA2 cTag-CLIP between cerebellar granule (vGluT1-Cre) and Purkinje cells (Pcp2-Cre). Scatter plot shows the correlation of read counts on CLIP peaks between NOVA2 cTag^vGlut1-Cre^-CLIP to NOVA2 cTag^Pcp2-Cre^-CLIP. Each *black* dot represents a comparable CLIP peak between two neuronal type. Each *magenta* or *orange* dot represents a significantly enriched CLIP peak in either NOVA2 cTag^vGlut1^-^Cre^-CLIP or cTag^Pcp2-Cre^-CLIP, respectively (FDR<0.05, |log2 FC|>=1). R: correlation coefficient. (D) Enrichment of YCAY motifs around NOVA2 cTag-CLIP peaks. (E) Manual dissection images of Purkinje cell layer from cerebellar slices of Control^tdTomato;Pcp2-Cre^ reporter mice. (F) Neuronal-type marker enrichment in and depletion from RNA-seq libraries prepared from dissected out Purkinje cell layer. Purkinje cell enriched markers (*orange*). Granule cell enriched markers (*magenta*). (G) Transcript abundance changes upon NOVA2 depletion from Purkinje cells determined by RNA-seq. Each *black* dot represents a transcript. Each *blue* or *red* dot represents a significantly decreased or increased transcript in the Purkinje cell layer of Nova2-cKO^Pcp2-Cre^ mouse, respectively (n=3, FDR<0.01, |log2FC|>=1, TPM average >= 2). (H) Venn diagram representing NOVA2 target AS events number overlapping or differentially regulated among three neuronal-linages. (I) AS changes between Purkinje cells and granule cells. Scatter plot shows the exon inclusion rate of Purkinje cells and granule cells determined by TRAP (Mellén et al., 2012). Each *grey* dot represents a comparative AS event. Each *black* dot indicates a significantly changed AS event (total 6,163 events) (n=4, FDR<0.05, |*Δ*I|>=0.1).

**Figure S4.**
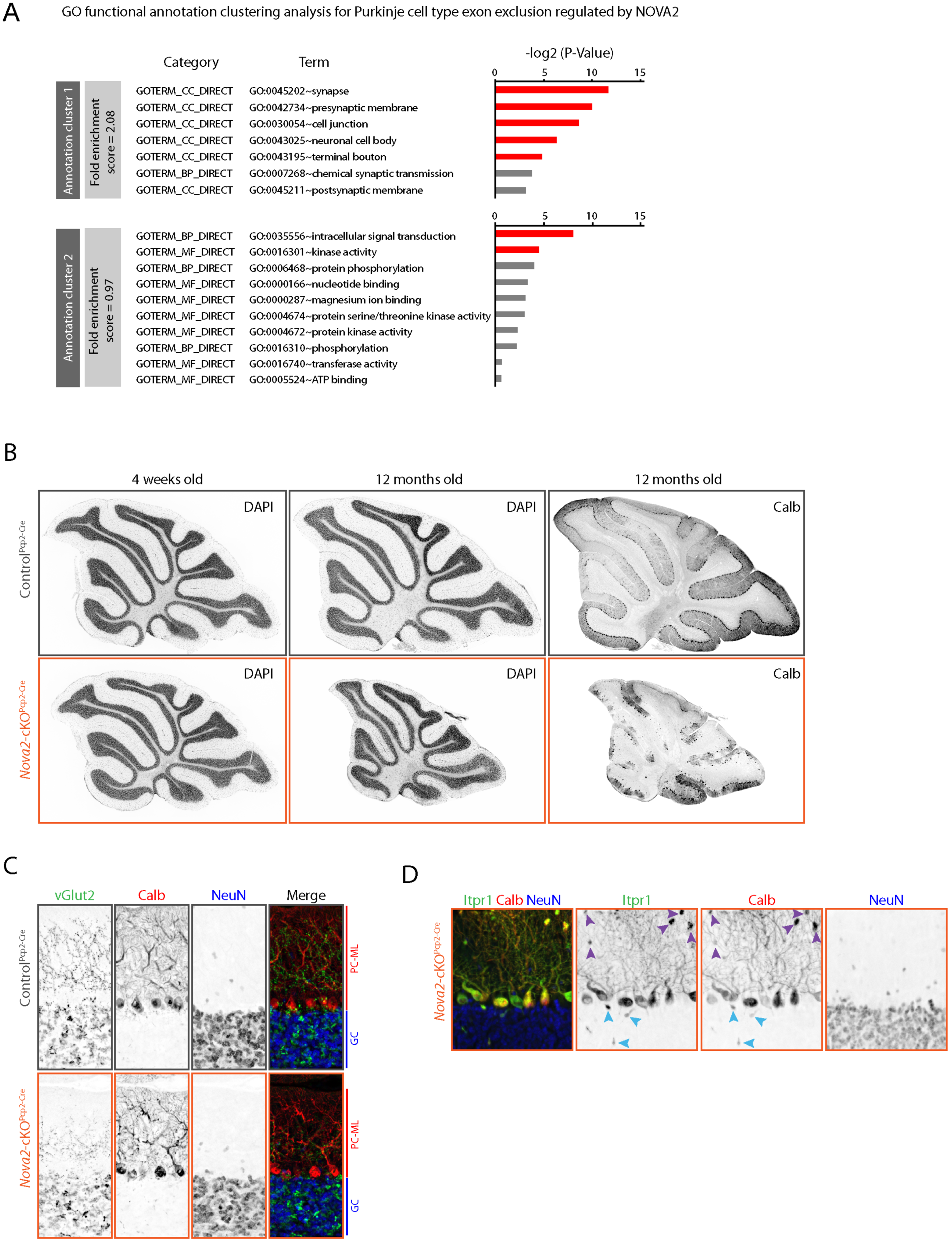
Related to Figure 4. Purkinje cell specific *Nova2* deficiency leads progressive motor discoordination and cerebellar atrophy. (A) GO functional annotation clustering analysis for the genes used in Figure 3H. *Red bars* represent the GO term having p<0.05. (B) Comparative granule cell layer thickness. DAPI staining images in 4 weeks and 12 months old Control^tdTomato; Pcp2-Cre^ and Nova2-cKO^tdTomato; Pcp2-Cre^ and Calbindin D28 immunostaining images at 12 months old. (C) The climbing fiber innervation defect and reduced Purkinje molecular layer thickness upon Purkinje cell specific *Nova2* deficiency. vGlut2 (*green*, climbing fiber terminal marker), calbindin D28 (*red*, Purkinje cells marker), and NeuN (*blue*, granule cell marker) immunofluorescent staining images in the 4 weeks old Control^Pcp2-Cre^ and *Nova2-* cKO^Pcp2-Cre^. (D) The swelling Purkinje cell’s neurites upon *Nova2* deficiency. ITPR1 (*green*), Caibindin D28 (*red*), and NeuN (*blue*) immunofluorescent staining images from 16 weeks old *Nova2*-cKO^Pcp2-Cre^ mouse. *Purple* and *blue* arrowheads represent swelling neurites in molecular layer or granule cell layer, respectively.

**Figure S5.**
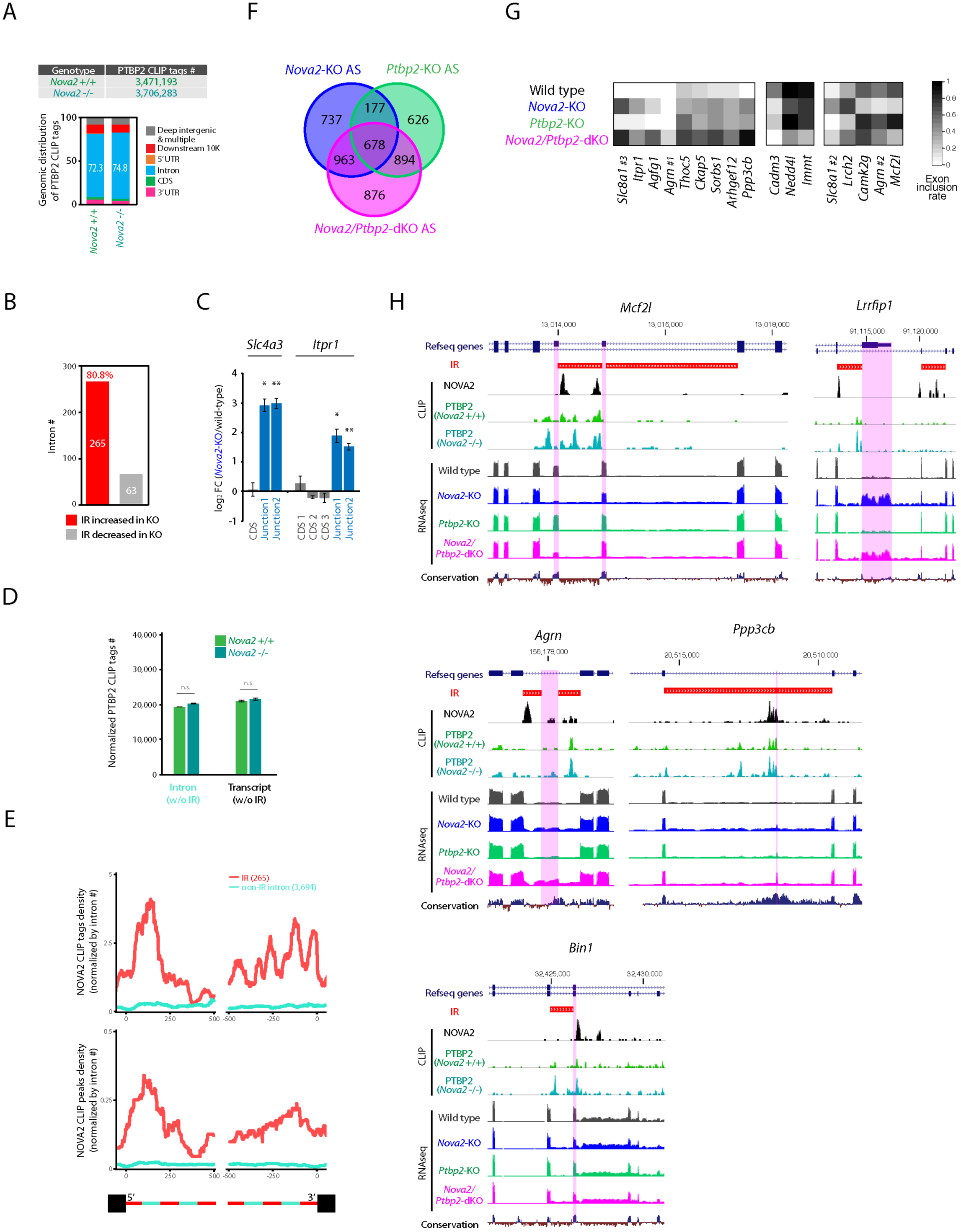
Related to Figure 5. NOVA2 regulates IR to serve as a *cis*-acting scaffold element for AS factor PTBP2. (A) PTBP2 CLIP results in wild type and *Nova2-*KO mouse. Total PTBP2 unique CLIP tag number from the e18.5 cortex of wild type and *Nova2-*KO mouse (*upper panel*) (three independent CLIP experiments used three biological replicates). Genomic distribution of PTBP2 unique CLIP tags (*lower graph*). (B) The number of NOVA2-dependent increased and decreased IR in *Nova2*-KO. (C) IR retaining exon-intron junction. qPCR quantification of *Slc4a3* and *Itpr1* RNA abundance with basic primer pairs designed both in coding sequence (CDS) and of RNA retaining exon-intron junction with primer pairs designed in either CDS or intron in e18.5 wild type and *Nova2*-KO mice (n=3, *; p<0.05, **; p<0.01). (D) PTBP2 CLIP tag number counted on all introns excepting IR, and full-length transcript excepting IR, which were normalized by total PTBP2 CLIP tag number of each replicate (n=3). Transcripts retaining IR or introns from these were subjected to analysis. (E) NOVA2 CLIP tags or peaks distribution on IR or non-retained intron. Normalized NOVA2 CLIP tag (upper) or peak (*lower*) density on IR or non-retained intron. (F) Venn diagram representing NOVA2, PTBP2, and NOVA2/PTBP2 target AS events number overlapping or differentially regulated among e18.5 cortex of *Nova2*-KO*, Ptbp2*-KO, and *Nova2/Ptbp2*-dKO. (G) Examples of AS changes in *Nova2-*KO, Ptbp2-KO, and *Nova2/Ptbp2*-dKO. Heatmaps represent the exon inclusion rate in either NOVA2 or PTBP2, or both absence condition *in vivo. Left* 9 columns show the additively or synergistically skipped by NOVA2 and PTBP2 in normal condition. *Middle* 3 columns show the additively or synergistically included by NOVA2 and PTBP2 in wild type. *Right* 5 columns show the attenuated-type AS events which are significantly changed in single-KO mouse but comparative in *Nova2/Ptbp2*-dKO with wild type. (H) UCSC genome browser views of the NOVA2-dependent IR containing transcript with PTBP2 CLIP peaks changes.

Table S1. Related to Figure 1–3, and 5. RBPs target AS events and NOVA2 target AS events in selective neuronal cell-types.

Table S2. Related to Figure 2, 3, and 5. Primers list.

Table S3. Related to Figure 1–3, and 5–6. NOVA2 cTag-CLIP and RNA-seq listed used in this study.

Movie S1. Related to Figure 4. Motor coordination defect in 13 weeks old Nova2-cKO^Pcp2-Cre^ Mouse.

